# Sotagliflozin Reprograms Hepatic Lipid Metabolism and Attenuates Atherosclerosis Beyond Empagliflozin in CKM Mice

**DOI:** 10.64898/2026.09.25.754502

**Authors:** Keith Saum, Jessica Sim, Lixia Zeng, Charlie Steward, Pradeep Kayampilly, Alethia Dixon, Anand Singh, Ahmed Abdel-Latif, Subramaniam Pennathur

**Affiliations:** Department of Internal Medicine, Division of Nephrology, University of Michigan, Ann Arbor, MI, USA; The University of Toledo College of Medicine and Life Sciences, Toledo, OH, USA; Department of Internal Medicine, Division of Cardiology, University of Michigan, Ann Arbor, MI, USA; Ann Arbor Veteran Affairs Healthcare System, Ann Arbor, MI, USA; Molecular and Integrative Physiology, University of Michigan, Ann Arbor, MI, USA

## Abstract

**Background:** Sodium-glucose cotransporter-2 (SGLT2) inhibitors reduce cardiovascular mortality and chronic kidney disease (CKD) progression in cardiovascular-kidney-metabolic (CKM) syndrome, but this benefit derives primarily from reductions in heart failure and cardiovascular mortality rather than in atherothrombotic events. Sotagliflozin, which inhibits both intestinal SGLT1 and renal SGLT2, is the only drug in this class shown to lower rates of myocardial infarction and stroke in CKD. However, its mechanism remains unclear, and its effectiveness in non-diabetic CKD has not been studied. We hypothesized that dual SGLT1/2 inhibition reprograms systemic substrate utilization toward fatty acid oxidation, which may partially account for this protective effect.

**Methods:** CKD was induced in male low-density lipoprotein receptor-deficient mice by 5/6 nephrectomy. Sham and CKD mice received a high-fat diet and drinking water containing sotagliflozin, empagliflozin, or vehicle for 16 weeks, at doses that produced equivalent fractional excretion of glucose. We quantified atherosclerosis *en face* and in the aortic sinus. Indirect calorimetry, echocardiography, hepatic RNA sequencing, plasma acylcarnitine profiling, and *in vivo* deuterium labeling of palmitate were performed.

**Results:** At matched levels of glucosuria and plasma total cholesterol, sotagliflozin decreased body mass, atherosclerosis burden, triglycerides, and TNF-α more than empagliflozin, and raised postprandial total GLP-1 approximately 3-fold, while both drugs similarly improved albuminuria and diastolic function. Sotagliflozin increased energy expenditure independent of body mass and lowered respiratory exchange ratio despite compensatory hyperphagia. Sotagliflozin reduced hepatic lipogenic genes and *de novo* lipogenic flux, while increasing β-oxidation and ketogenesis; empagliflozin only activated the oxidative pathway.

**Conclusions:** In experimental CKM syndrome, dual SGLT1/2 inhibition reduces atherosclerosis and metabolic syndrome more effectively than selective SGLT2 inhibition and is associated with reprogramming of hepatic lipid metabolism. These results suggest that sotagliflozin could be an attractive candidate for mitigating atherosclerotic risk in non-diabetic CKD.

**Graphical Abstract:** 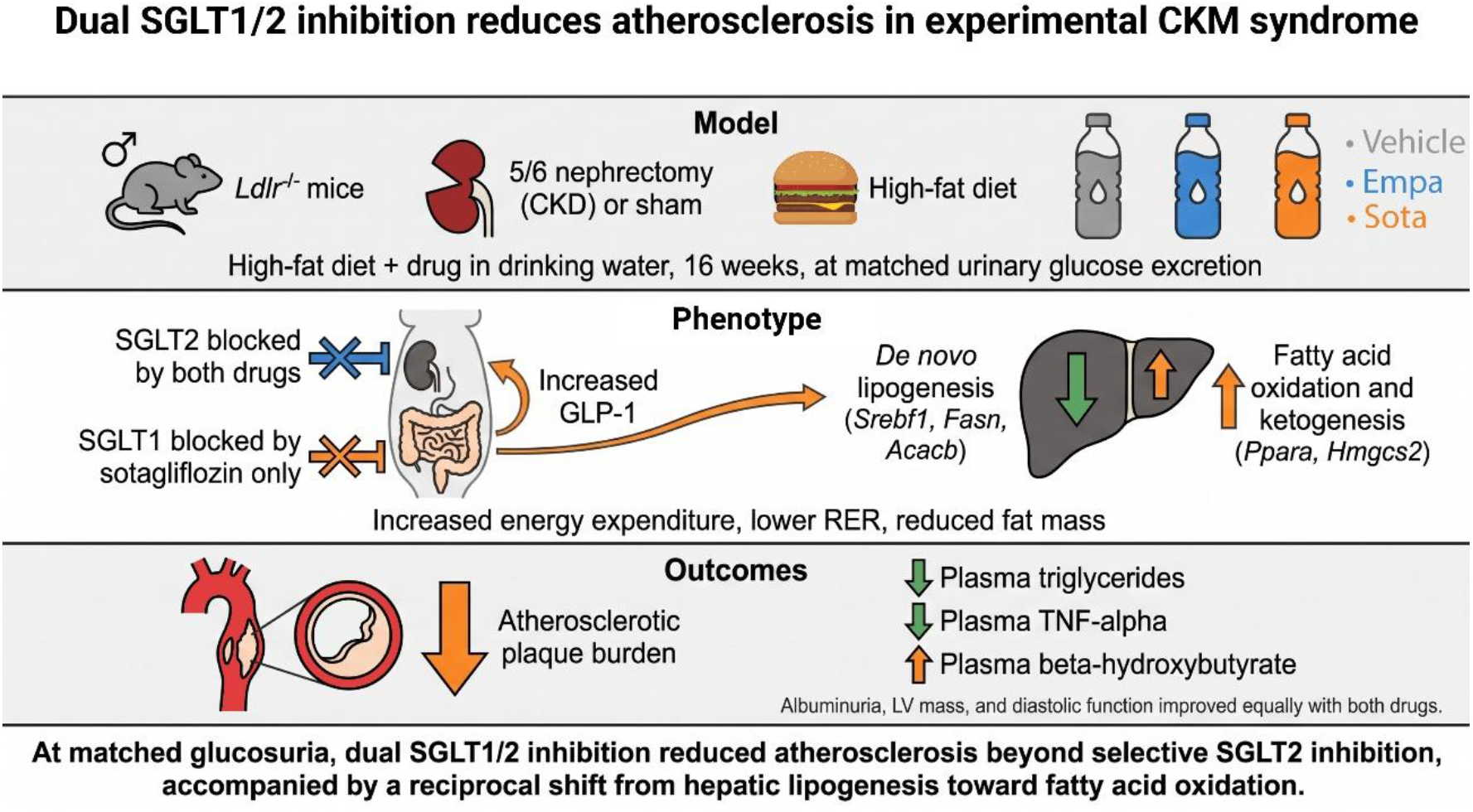

## Introduction

Patients with chronic kidney disease (CKD) face an increased risk of cardiovascular disease (CVD).^1^ As a result, CVD remains the leading cause of death among CKD patients and poses a greater risk than progression to kidney failure.^2^ This link between CVD and CKD is now emphasized through the concept of cardiovascular-kidney-metabolic (CKM) syndrome, which also incorporates metabolic risk factors and their combined impact on morbidity and mortality.^3^ Recent data indicate that most US adults now meet criteria for CKM stages 2-4, which include established CKD or metabolic risk factors at one end and clinically manifested CVD at the other.^4^ Consequently, kidney function has been integrated into standard risk assessment models, with the latest AHA PREVENT equations incorporating estimated glomerular filtration rate and urinary albumin-to-creatinine ratio.^5^ Therefore, effective strategies to prevent and treat CVD in this high-risk population remain an unmet need.

Sodium-glucose cotransporter-2 (SGLT2) inhibitors are now a foundational therapy across the spectrum of CKM syndrome.^6^ Large-scale trials have demonstrated their robust efficacy in reducing cardiovascular mortality in patients with type 2 diabetes mellitus, heart failure, and CKD.^7^ However, this cardiovascular benefit is driven primarily by reductions in cardiovascular death and heart failure hospitalizations, with minimal impact on myocardial infarction or stroke. In contrast, glucagon-like peptide-1 (GLP-1) receptor agonists demonstrate consistent reductions in myocardial infarction and stroke in patients with and without diabetes.^8^ Furthermore, adding an SGLT2 inhibitor to a GLP-1 receptor agonist does not reduce cardiovascular risk beyond what either agent achieves alone.^9^ These findings indicate that selective SGLT2 inhibition does not fully address the significant atherothrombotic risk in CKD, and that incretin signaling plays a role in vascular protection.

Sotagliflozin is unique among SGLT2 inhibitors in that it inhibits both SGLT2 and SGLT1. SGLT2 is primarily expressed in the proximal tubule and accounts for about 90% of renal glucose reabsorption, whereas SGLT1 is highly expressed in the small intestine, where it facilitates absorption of dietary glucose and galactose and also contributes to the remaining 10% of renal glucose reabsorption.^10^ Inhibition of intestinal SGLT1 by sotagliflozin reduces postprandial glucose excursion and may offer additional cardiometabolic benefits beyond SGLT2 inhibition alone. When compared to selective SGLT2 inhibition by empagliflozin, dual inhibition of SGLT1/2 by sotagliflozin produces greater reductions in postprandial glucose and triglycerides along with greater elevations in postprandial GLP-1 levels.^11^ Furthermore, loss-of-function variants in *SLC5A1*, the gene encoding SGLT1, are associated with a lower incidence of heart failure, diabetes, and cardiovascular death.^12^ Clinical evidence from the SCORED trial demonstrated that sotagliflozin significantly reduces major adverse cardiovascular events, including both myocardial infarction and stroke, in patients with CKD and diabetes.^13,14^ However, the SCORED trial only enrolled CKD patients with type 2 diabetes and was stopped early because of a lack of funding, which limited the number of events and power to assess changes in CKD progression. It is unknown if the atherothrombotic benefits of sotagliflozin extend to the non-diabetic CKD population, and the specific contributions of dual SGLT1/2 inhibition to atherosclerosis development, metabolism, and inflammation remain unknown.

Given the frequent coexistence of CKD and metabolic dysfunction in CVD, we used a pathophysiological murine model combining 5/6 nephrectomy to induce CKD in Ldlr^-/-^ mice with a high-fat diet to induce metabolic syndrome and atherosclerosis.^15–18^ This model recapitulates key features of CKM syndrome, including albuminuria, diastolic dysfunction, systemic inflammation, and insulin resistance. We hypothesized that treatment with the dual SGLT1/2 inhibitor sotagliflozin would reduce systemic glucose metabolism and shift systemic substrate utilization toward fatty acid oxidation, thereby attenuating atherosclerosis and metabolic syndrome compared with selective SGLT2 inhibition by empagliflozin. Using indirect calorimetry, metabolic profiling, and *in vivo* lipogenic flux analysis, we demonstrate that reductions in atherosclerosis and metabolic syndrome with sotagliflozin are accompanied by a systemic shift toward fatty acid oxidation, lower hepatic *de novo* lipogenesis, and higher endogenous GLP-1, effects not seen with selective SGLT2 inhibition. These findings identify metabolic reprogramming downstream of intestinal SGLT1 inhibition as a candidate mechanism underlying the atherothrombotic protection conferred by sotagliflozin and support further evaluation in non-diabetic CKD.

## Methods

### Animals

All animal experiments were performed with the approval of the University of Michigan Institutional Animal Care and Use Committee (IACUC). At 6 weeks of age, male *Ldlr*^−/−^ mice (B6.129S7-Ldlr^tm1Her/J^, The Jackson Laboratory) were subjected to either sham operation or 5/6 nephrectomy to induce CKD, as previously described.^15^ Briefly, the right kidney was removed in an initial procedure, followed by approximately two-thirds of the left kidney one week later. At 10 weeks of age, mice were transitioned to a high-fat diet (HFD; 20% protein, 21% fat with 0.21% cholesterol, and 50% carbohydrates by weight; D12079B, Research Diets Inc.) to induce hypercholesterolemia and block randomized to receive sotagliflozin (SOTA, 30mg/kg/day), empagliflozin (EMPA, 20mg/kg/day), or vehicle continuously via drinking water (n=20 per treatment arm within each surgical group, 2-3 mice/cage). SOTA and EMPA (Med Chem Express, Cat. #s HY-15516 and HY-15409) were dissolved in 100% ethanol with sonication, then diluted in water containing 0.25% Tween-80 to a final ethanol concentration of 1%. Vehicle-treated mice received drinking water containing 1% ethanol and 0.25% Tween-80. We selected final SOTA and EMPA concentrations in a preliminary dose-finding cohort, in which we calculated fractional glucose excretion from paired plasma and urine glucose and creatinine measurements after 7 days of treatment. Blood and urine glucose levels were measured using a handheld glucometer (AimStrip Plus, Germaine Laboratories Inc.) and an enzymatic assay (Abcam cat#ab65333), respectively. Plasma creatinine was measured by highly specific electrospray ionization and tandem LC-MS, as previously described.^18^ The same measurement was repeated in the main cohort to confirm sustained pharmacodynamic activity and comparable renal SGLT2 inhibition between groups. All mice received HFD and drinking water ad libitum for 16 weeks before blood and tissue collection. We refreshed drinking water every 48 hours and measured body weight and food intake weekly. Because human studies indicate that cardiorenal protection by SGLT2 inhibitors is independent of sex, and female mice are less susceptible to diet-induced atherosclerosis than males, we used only male mice in this study.^7,13,19^ Detailed methods are provided in the Supplemental Material.

### Statistics

Data are presented as mean ± SEM. Normality was assessed by the Shapiro-Wilk. Variables that deviated from normality were log-transformed before analysis or compared by nonparametric tests, as indicated in the corresponding figure legend. We analyzed measurements from both surgical groups using two-way ANOVA with surgical group (sham or CKD) and treatment (vehicle, EMPA, or SOTA) as fixed factors, followed by Tukey’s multiple comparisons test. Measurements obtained only in CKD animals were analyzed by one-way ANOVA with treatment group as a fixed factor. Animals that died before the study ended were excluded from terminal endpoints. Serial measurements, including body weight, blood glucose, and total GLP-1 during OGTT, were analyzed by two-way repeated-measures ANOVA. Glycemic and incretin excursions were additionally summarized as total AUC by the trapezoidal method and compared as above. A two-sided *P*<0.05 was considered statistically significant. Analyses were performed using GraphPad Prism version 11 (GraphPad Software) and R version 4.6.1 (R Foundation for Statistical Computing).

## Results

### Dual SGLT1/2 inhibition attenuates diet-induced obesity and metabolic syndrome independent of kidney function

To determine whether dual SGLT1/2 inhibition confers cardiometabolic benefit beyond selective SGLT2 inhibition in CKM syndrome, male *Ldlr*^−/−^ mice underwent 5/6 nephrectomy or sham operation, were placed on a HFD, and were randomized to sotagliflozin (30 mg/kg per day), empagliflozin (20 mg/kg per day), or vehicle delivered continuously in drinking water for 16 weeks (Figure 1A). Sotagliflozin attenuated diet-induced weight gain in both sham and CKD groups, with divergence beginning at week 4 and widening progressively through study end (Figure 1B and 1C). Body weight in empagliflozin-treated mice did not differ from vehicle-treated mice at any time point in either surgical group. Despite these differences in body weight, sotagliflozin and empagliflozin lowered plasma creatinine and albuminuria to a similar degree compared to vehicle in CKD mice (Figure 1D and 1E) but did not change plasma Cystatin C levels (Figure S1E). Because plasma creatinine fell to a similar degree with both agents while only sotagliflozin reduced body mass, this change is unlikely to reflect differences in muscle mass. Fractional excretion of glucose was negligible with vehicle and increased similarly with sotagliflozin and empagliflozin in both surgical groups (Figure 1F), confirming comparable renal SGLT2 inhibition at the doses used. No differences were observed in serum electrolytes or hemoglobin between treatment groups in sham or CKD mice (Figure S1). Serum bicarbonate and anion gap did not differ between treatment groups in either surgical group (Figure S1F and S1G), indicating that neither agent worsened CKD-induced metabolic acidosis.

**Figure 1.**
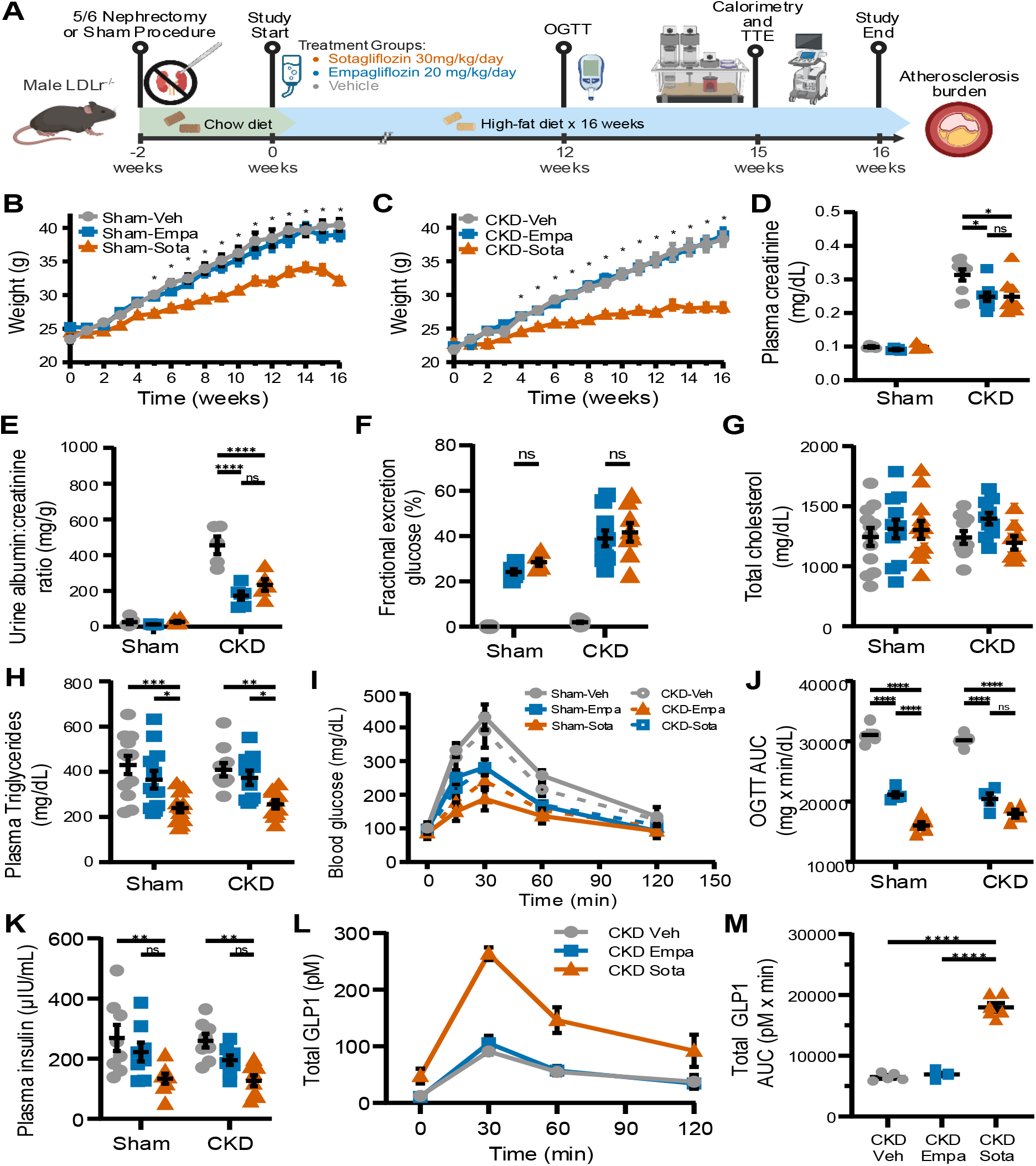
Dual SGLT1/2 inhibition with sotagliflozin attenuates diet-induced obesity and metabolic syndrome independent of kidney function. (A) Male *Ldlr*^−/−^ mice underwent sham operation or 5/6 nephrectomy, were transitioned to a HFD at 10 weeks of age, and were randomized to vehicle, EMPA (20 mg/kg/day), or SOTA (30 mg/kg/day) in drinking water for 16 weeks. (B-C) Body weight over the treatment period in sham (B) and CKD (C) mice. Asterisks represent P<0.05 between EMPA and SOTA mice. (D) Plasma creatinine. (E) UACR. (F) Fractional excretion of glucose. (G) Total plasma cholesterol. (H) Plasma triglycerides. (I-J) Blood glucose during OGTT performed after 12 weeks of treatment (I) and corresponding AUC (J) (n=5/group). (K) Plasma insulin. (L-M) Plasma total GLP-1 during OGTT in CKD mice (L) and corresponding AUC (M) (n=5-6/group). Data are shown as mean±SEM. Body weight and time-course data were analyzed by two-way repeated-measures ANOVA and all other comparisons by two-way ANOVA, each with Tukey’s multiple comparisons test. ✱*P*<0.05, ✱✱*P*<0.01, ✱✱✱*P*<0.001, and ✱✱✱✱*P*<0.0001; ns, not significant. AUC, area under the curve; CKD, chronic kidney disease; EMPA, empagliflozin; GLP-1, glucagon-like peptide-1; HFD, high-fat diet; OGTT, oral glucose tolerance test; SOTA, sotagliflozin; and UACR, urine albumin-to-creatinine ratio.

Sotagliflozin also attenuated features of metabolic syndrome more than empagliflozin. Although plasma total cholesterol was unchanged by either treatment (Figure 1G), sotagliflozin lowered plasma triglycerides relative to both vehicle and empagliflozin in sham and CKD mice (Figure 1H). Oral glucose tolerance testing after 12 weeks of treatment revealed graded improvement in glycemic excursion across the three treatment arms (Figure 1I and 1J). Both inhibitors reduced glycemic excursion relative to vehicle, and sotagliflozin produced a further reduction beyond empagliflozin in sham mice, with a similar but nonsignificant difference in CKD mice. Plasma insulin was lower with sotagliflozin than with vehicle in both surgical groups, whereas empagliflozin was indistinguishable from vehicle (Figure 1K), consistent with improved insulin sensitivity. Sotagliflozin also amplified the postprandial incretin response. In CKD mice, total GLP-1 rose threefold above vehicle and empagliflozin at 30 minutes after oral glucose and remained elevated through 120 minutes, yielding a markedly greater GLP-1 area under the curve (Figure 1L and 1M). Empagliflozin did not differ from vehicle. Thus, at matched renal SGLT2 inhibition, only dual SGLT1/2 inhibition limited weight gain, reduced triglycerides and insulin, and increased endogenous GLP-1, without altering cholesterol exposure.

### Dual SGLT1/2 inhibition reduces atherosclerosis beyond selective SGLT2 inhibition

We next quantified atherosclerotic burden after 16 weeks of treatment. Vehicle-treated CKD mice developed greater lesion burden than sham controls in both the whole aorta and the aortic sinus, confirming that reduced kidney function accelerated atherogenesis in this CKM model (Figure 2A through 2D, *en face* effect for CKD, F(1, 66) = 15.84, p=0.0002, η²_p_ = .19). Both SGLT inhibitors reduced lesion burden in sham and CKD mice, but sotagliflozin reduced plaque burden area to a greater extent than empagliflozin in both surgical groups (Figure 2C and 2D, *en face* effect for treatment, F(2, 66) = 20.92, p <0.0001, η²_p_ = .39). Notably, this graded protection occurred without any difference in total plasma cholesterol between treatment groups (Figure 1G), indicating that the additional benefit of dual SGLT1/2 inhibition was not mediated by reduced cholesterol exposure.

**Figure 2.**
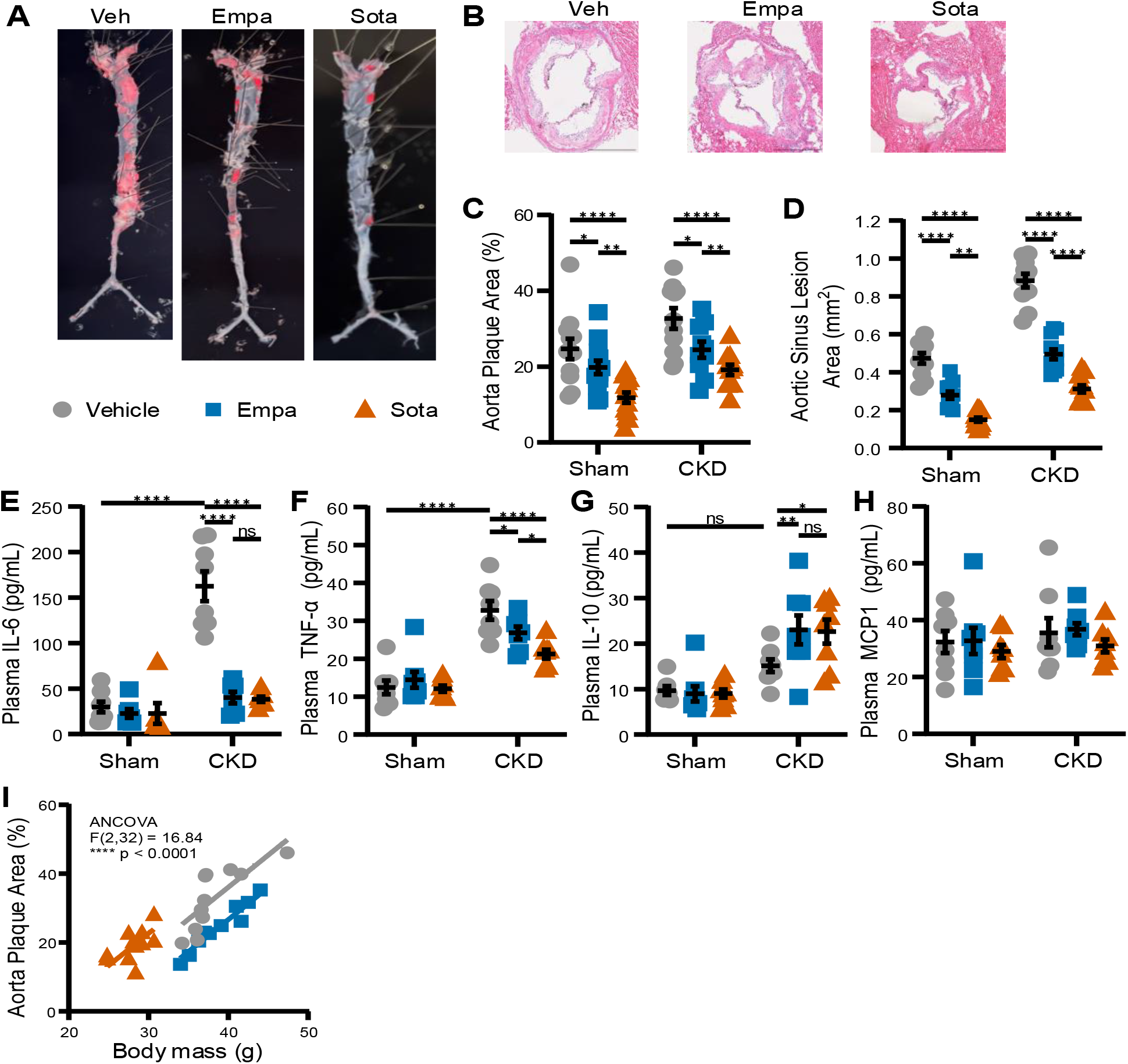
Sotagliflozin reduces atherosclerotic burden and systemic inflammation beyond selective SGLT2 inhibition. Atherosclerotic burden was quantified after 16 weeks of treatment in sham and CKD mice (n=12/group), and plasma inflammatory mediators were measured by multiplex immunoassay. (A) Representative *en face* preparations of Oil Red O-stained aortas. (B) Representative H&E-stained aortic sinus sections. Scale bar, 500 µm. (C) *En face* aortic plaque area, expressed as percent of total aortic surface area. (D) Aortic sinus lesion area summed across serial sections. (E-H) Plasma concentrations of IL-6 (E), TNF-α (F), IL-10 (G), and MCP-1 (H). All lesion quantification was verified by 2 blinded observers. (I) ANCOVA of *en face* aortic plaque area against total body mass in CKD mice, with treatment as a fixed factor and body mass as the covariate. Data are shown as mean±SEM. Comparisons were analyzed by two-way ANOVA with Tukey’s multiple comparisons test. ✱*P*<0.05, ✱✱*P*<0.01, ✱✱✱*P*<0.001, and ✱✱✱✱*P*<0.0001; ns, not significant. CKD, chronic kidney disease; IL-6, interleukin 6; IL-10, interleukin 10; MCP-1, monocyte chemoattractant protein-1; and TNF-α, tumor necrosis factor alpha.

Because atherosclerosis in CKD is driven in part by systemic inflammation, we measured circulating inflammatory mediators. CKD raised plasma IL-6 and TNF-α relative to sham operation (Figure 2E and 2F). Both inhibitors reduced IL-6 to concentrations indistinguishable from those in sham mice, with no difference between agents (Figure 2E). TNF-α was likewise reduced by both drugs, although sotagliflozin lowered it further than empagliflozin (Figure 2F). The anti-inflammatory cytokine IL-10 rose with both treatments in CKD mice to a comparable degree (Figure 2G), whereas MCP-1 was unaffected by CKD or by either treatment (Figure 2H). Because adipose tissue is itself a source of circulating inflammatory mediators, and because sotagliflozin-treated mice weighed substantially less than vehicle- and empagliflozin-treated mice, both the reduction in plaque and the further reduction in TNF-α could in principle reflect lower adiposity alone. Aortic plaque area varied with body mass within each treatment group, yet treatment remained a significant determinant of lesion burden after adjustment for body mass by ANCOVA (Figure 2I, treatment effect, F(2, 32) = 16.84, p<0.0001, η²_p_ = .55), indicating that reduced adiposity does not fully account for the atheroprotection conferred by dual SGLT1/2 inhibition. Together, these findings establish that dual SGLT1/2 inhibition confers greater protection against atherosclerosis than selective SGLT2 inhibition in mice with CKM syndrome, and that this incremental benefit is accompanied by a further reduction in systemic TNF-α rather than by broad suppression of all inflammatory mediators.

### SGLT1 inhibition does not augment protection against CKD-induced diastolic dysfunction

To determine whether the metabolic divergence between sotagliflozin and empagliflozin extended to cardiac structure and function, we performed transthoracic echocardiography after 15 weeks of treatment. Heart rate was comparable across treatment groups (Figure 3A), and systolic function was preserved in both sham and CKD mice with similar ejection fraction and fractional shortening (Figure 3B and 3C). CKD induced cardiac hypertrophy with impaired diastolic function. Compared with sham mice, vehicle-treated CKD mice had greater corrected left ventricular mass (Figure 3D) and a higher heart weight-to-tibia length ratio (Figure 3H), accompanied by elevated E/e′ and E/A ratios and isovolumic relaxation time (Figure 3E through 3G). The combination of preserved ejection fraction, ventricular hypertrophy, and impaired relaxation is consistent with the heart failure with preserved ejection fraction phenotype, common in CKM syndrome. In CKD mice, sotagliflozin and empagliflozin each lowered corrected left ventricular mass and heart weight-to-tibia length ratio toward sham values (Figure 3D and 3H) and normalized E/e′ ratio, E/A ratio, and isovolumic relaxation time (Figure 3E through 3G). No measure of cardiac structure or diastolic function differed between the two agents, and neither agent altered any parameter in sham mice.

**Figure 3.**
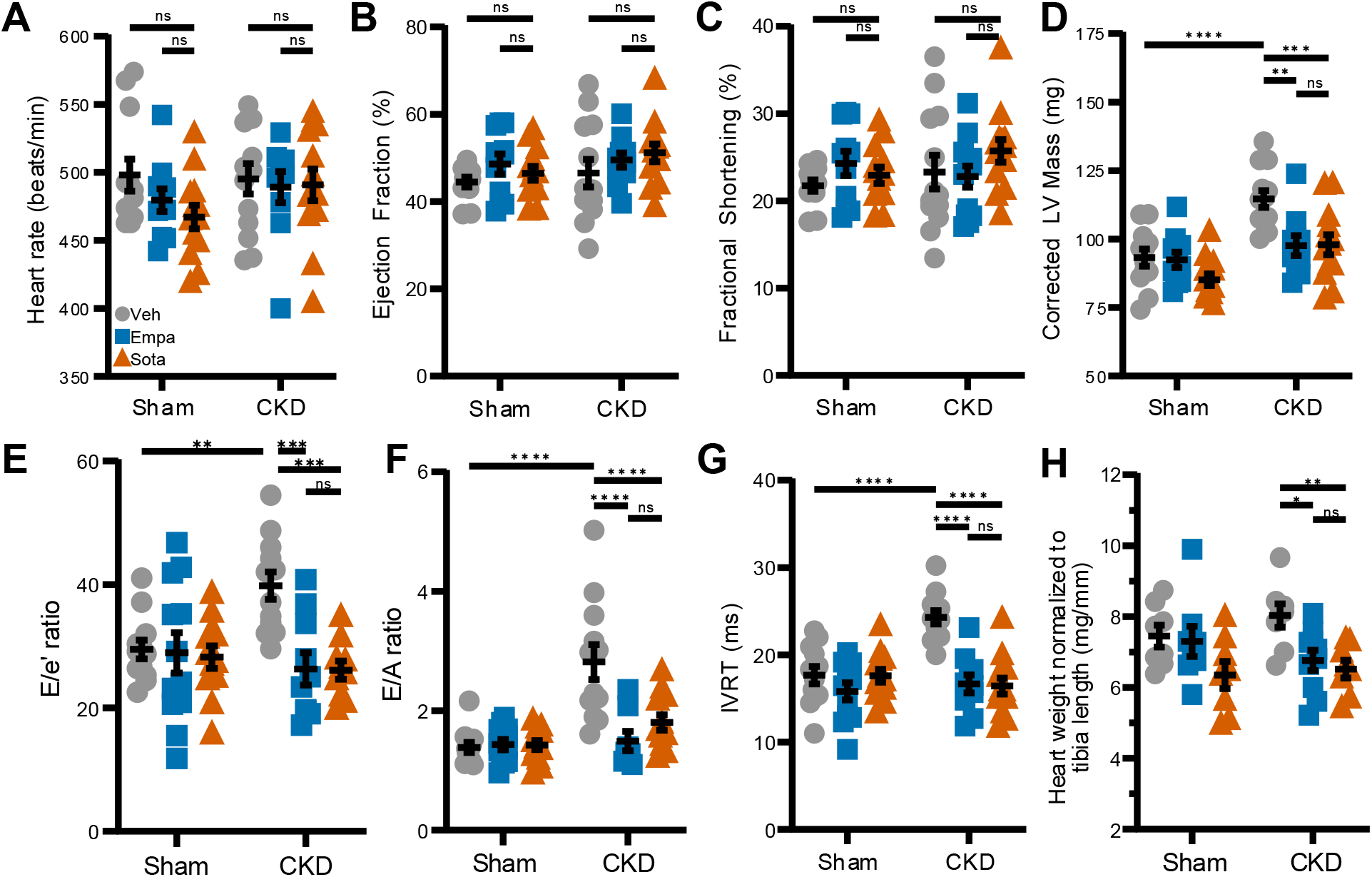
SGLT1 inhibition does not augment protection against CKD-induced diastolic dysfunction. Transthoracic echocardiography was performed in a subset of sham and CKD mice after 15 weeks of treatment (n=10-12/group). All parameters were measured at least three times and averaged for analysis. (A) Heart rate. (B) Ejection fraction. (C) Fractional shortening. (D) Corrected LV mass. (E) E/e′ ratio. (F) E/A ratio. (G) IVRT. (H) Heart weight normalized to tibia length. Data are shown as mean±SEM. Comparisons were analyzed by two-way ANOVA with Tukey multiple comparisons test. ✱*P*<0.05, ✱✱*P*<0.01, ✱✱✱*P*<0.001, and ✱✱✱✱*P*<0.0001; ns, not significant. CKD, chronic kidney disease; E/A, ratio of early to late mitral inflow velocity; E/e′, ratio of early mitral inflow velocity to early diastolic mitral annular velocity; IVRT, isovolumic relaxation time; and LV, left ventricle.

### Sotagliflozin increases energy expenditure and shifts substrate utilization toward fatty acid oxidation

To identify the basis of the divergent weight phenotype, we performed indirect calorimetry after 14 weeks of treatment. Body composition analysis showed that the reduced body mass in sotagliflozin-treated CKD mice was due solely to decreased adiposity, with fat mass significantly lower than in vehicle-treated mice (Figure 4A). Lean mass remained consistent across all three groups. Empagliflozin-treated mice did not differ from vehicle-treated mice in either compartment. Sotagliflozin increased energy expenditure throughout the 72-hour calorimetry monitoring, during both dark and light phases, whereas empagliflozin was indistinguishable from vehicle (Figure 4B through 4D). Both SGLT inhibitors increased dark-phase energy intake relative to vehicle, consistent with compensatory hyperphagia in response to urinary caloric loss (Figure 4E through 4G). Despite this greater intake, only sotagliflozin-treated mice maintained a neutral-to-negative apparent energy balance over the recording period, whereas vehicle- and empagliflozin-treated mice remained in positive balance (Figure 4H and 4I). To account for differences in body mass between groups, energy expenditure was reanalyzed using ANCOVA with total body mass as the covariate, which confirmed that expenditure remained higher with sotagliflozin, independent of total body mass (Figure 4J, ANCOVA effect for treatment, F(2, 12) = 8.76, p = 0.0045, η²_p_ = .59). Ambulatory activity also did not differ between groups (Figure 4K), excluding a locomotor contribution.

**Figure 4.**
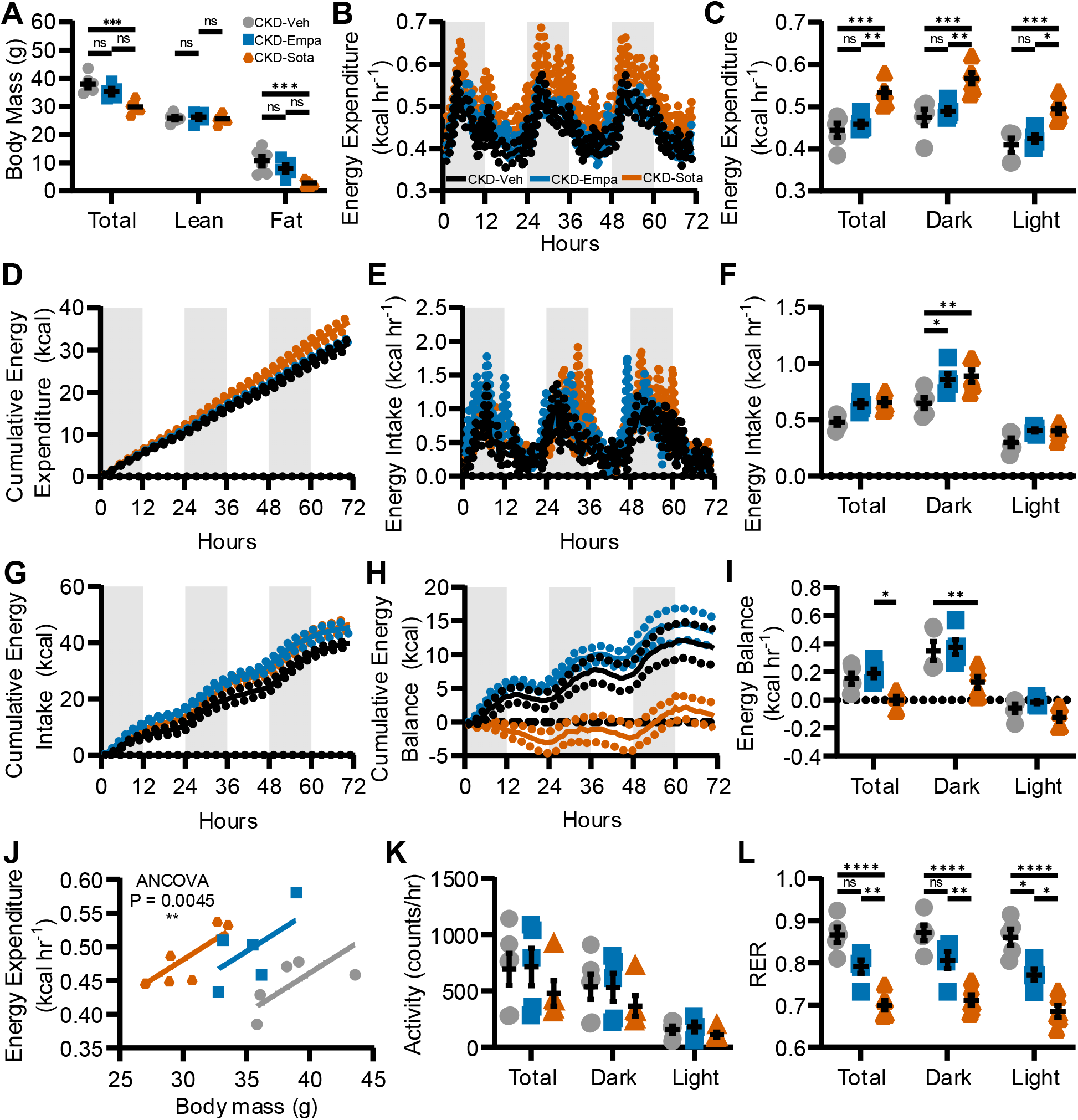
Sotagliflozin increases energy expenditure and shifts substrate utilization toward fatty acid oxidation. CKD mice underwent indirect calorimetry for 72 hours after 14 weeks of treatment (n=5/group), with body composition measured by EchoMRI immediately before and after the recording. (A) Total, lean, and fat mass. (B-C) Energy expenditure over the 72-hour recording (B) and averaged across the total recording and by dark and light phase (C). (D) Cumulative energy expenditure. (E-F) Energy intake over the recording period (E) and averaged across the total recording and by phase (F). (G) Cumulative energy intake. (H-I) Cumulative energy balance (H) and energy balance averaged across the total recording and by phase (I). (J) ANCOVA of energy expenditure against total body mass. (K) Ambulatory activity. (L) RER, calculated as VCO_2_/VO_2_. Shaded regions in B, D, E, G, and H denote the dark phase; solid lines indicate group means and dotted lines indicate SEM. Data are shown as mean±SEM. Energy expenditure was analyzed by ANCOVA against total body mass in CalR and presented per IICCC guidelines; all other comparisons were analyzed by one-way ANOVA with Tukey’s multiple comparisons test. ✱*P*<0.05, ✱✱*P*<0.01, ✱✱✱*P*<0.001, and ✱✱✱✱*P*<0.0001; ns, not significant; CKD, chronic kidney disease; IICCC, International Indirect Calorimetry Consensus Committee; RER, respiratory exchange ratio; VCO_2_, carbon dioxide production; and VO_2_, oxygen consumption.

The higher energy expenditure with sotagliflozin was accompanied by a shift in the fuel oxidized to sustain it. The respiratory exchange ratio was substantially lower in sotagliflozin-treated mice during both dark and light phases, approaching values indicative of near-exclusive fatty acid oxidation, whereas empagliflozin produced only a modest reduction confined to the light phase (Figure 4L). Sham mice displayed the same calorimetric phenotype (Figure S2), indicating that these effects of dual SGLT1/2 inhibition are independent of reduced kidney function. Taken together, sotagliflozin produced a coordinated increase in energy expenditure and a shift toward fat as the predominant oxidative substrate, sufficient to offset compensatory hyperphagia and drive selective loss of fat mass.

### Sotagliflozin reprograms hepatic lipid metabolism away from *de novo* lipogenesis and toward fatty acid oxidation

To define the transcriptional basis for this metabolic shift, we performed RNA sequencing on liver tissue from vehicle-, empagliflozin-, and sotagliflozin-treated CKD mice. Principal component analysis separated vehicle-treated mice from both treatment groups along the first principal component, with sotagliflozin-treated mice displaced furthest from vehicle and only partially overlapping the empagliflozin group (Figure 5A). Sotagliflozin altered a larger set of hepatic transcripts relative to vehicle than did empagliflozin (Figure 5B and 5C), and the empagliflozin response was largely contained within that of sotagliflozin: 107 of the 162 genes altered by empagliflozin relative to vehicle (66%) were also altered by sotagliflozin, while the sotagliflozin response comprised 283 genes, 176 of which were not shared with empagliflozin (Figure 5E). Direct comparison of the two drugs revealed a coherent lipogenic signature, with 15 genes uniquely altered by sotagliflozin relative to vehicle or empagliflozin (supplemental table S4). Sotagliflozin uniquely suppressed *Fasn*, *Acacb*, *Sqle*, and *Cyp7a1* relative to empagliflozin, while inducing transcripts associated with lipid-handling macrophages and stress response, including *Trem2*, *Gpnmb*, *Lgals1*, *Mmp12*, *Hmox1*, and *Atf3* (Figure 5D).

**Figure 5.**
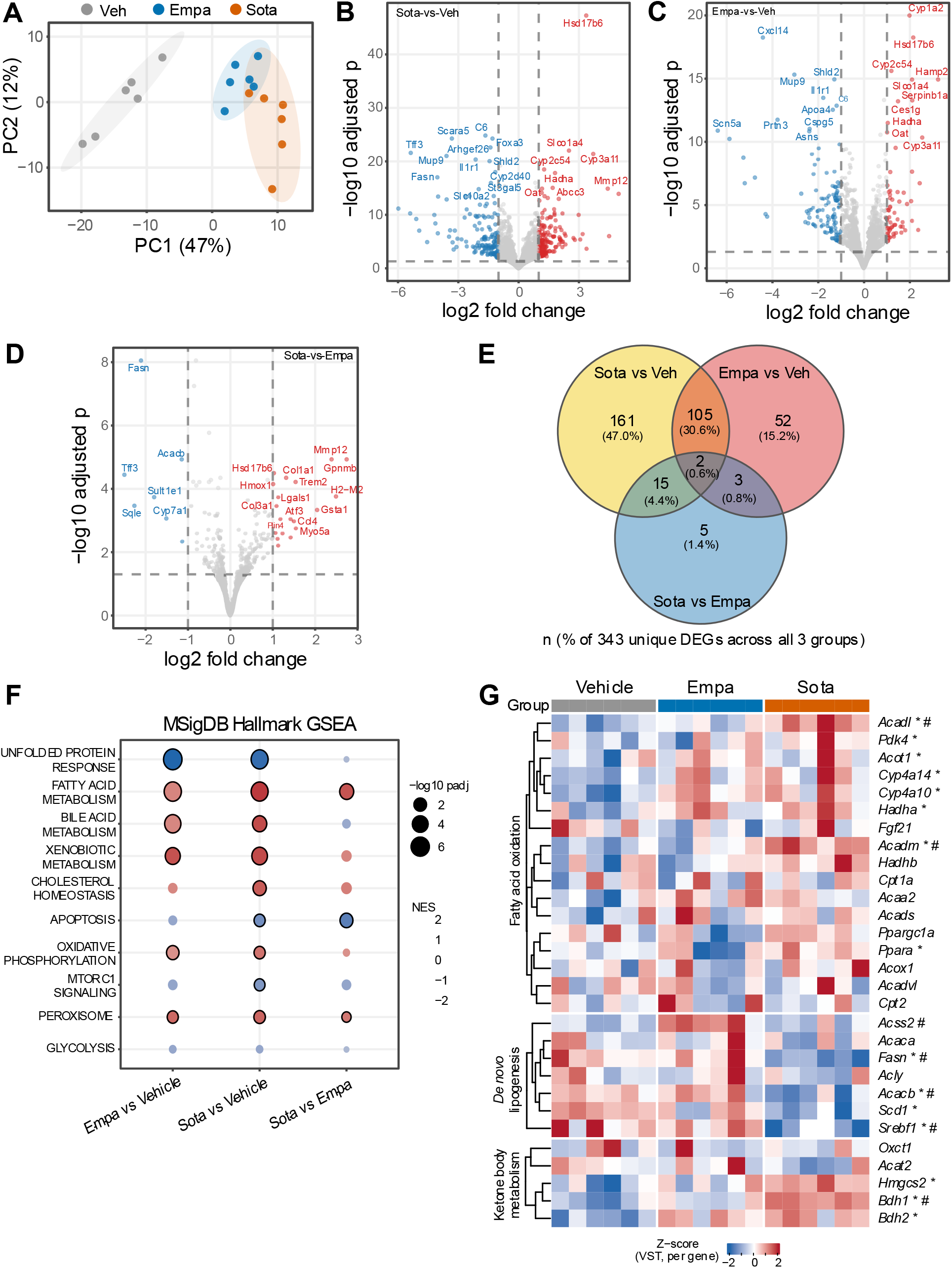
Sotagliflozin reprograms hepatic lipid metabolism away from *de novo* lipogenesis and toward fatty acid oxidation. Bulk RNA sequencing was performed on liver tissue from vehicle-, EMPA-, and SOTA-treated CKD mice (n=6/group). Differential expression was assessed with DESeq2 using Benjamini-Hochberg correction, with significance defined as FDR<0.05. (A) PCA of variance-stabilized transformed counts; ellipses denote 95% CIs. (B-D) Volcano plots of differential expression for SOTA versus vehicle (B), EMPA versus vehicle (C), and SOTA versus EMPA (D). Vertical dashed lines denote log_2_ fold change of ±1 and horizontal dashed lines an adjusted *P* of 0.05. Genes meeting both thresholds are colored red for upregulated and blue for downregulated. (E) Venn diagram of differentially expressed genes (DEGs, FDR<0.05 and |log2FC|>1) across the three pairwise comparisons, expressed as counts and as a percent of the 343 unique DEGs identified across all comparisons. (F) GSEA against the MSigDB Hallmark collection. Dot color denotes normalized enrichment score, dot size denotes −log_10_ adjusted *P*, and black outlines denote gene sets reaching an adjusted *P*<0.05. (G) Heatmap of genes involved in fatty acid oxidation, *de novo* lipogenesis, and ketone body metabolism, displayed as per-gene z-scores of variance-stabilized transformed counts with rows ordered by hierarchical clustering within each pathway. In G, ✱ indicates adjusted *P*<0.05 versus vehicle; and #, adjusted *P*<0.05 versus EMPA. Each column represents an individual mouse. CKD, chronic kidney disease; EMPA, empagliflozin; and SOTA, sotagliflozin.

Gene set enrichment analysis against pathways in the MSigDB Hallmark collection showed that both agents enriched fatty acid metabolism, peroxisome, bile acid metabolism, and xenobiotic metabolism relative to vehicle (Figure 5F). Critically, fatty acid metabolism and peroxisome gene sets remained significantly enriched in the direct sotagliflozin versus empagliflozin comparison, establishing that dual SGLT1/2 inhibition engaged these programs more strongly than selective SGLT2 inhibition, rather than merely reproducing them. Beyond lipid metabolism, both agents suppressed the unfolded protein response relative to vehicle, and sotagliflozin additionally suppressed the apoptosis gene set relative to empagliflozin (Figure 5F), consistent with reduced hepatic ER stress accompanying the lower lipogenic and lipid-storage burden.

Examination of individual lipid-handling genes revealed reciprocal regulation of synthesis and oxidation (Figure 5G). Sotagliflozin suppressed the *de novo* lipogenic program, including the master regulator *Srebf1* and its targets *Fasn*, *Acacb*, and *Scd1*, with *Fasn*, *Acacb*, and *Srebf1* also reduced relative to empagliflozin. Concurrently, sotagliflozin induced the transcriptional regulator of fatty acid oxidation, *Ppara*, along with downstream targets of β- and ω-oxidation, including *Acadl*, *Acadm*, *Acot1*, *Hadha*, *Cyp4a10*, *Cyp4a14*, and *Pdk4*, and upregulated the ketogenic enzymes *Hmgcs2*, *Bdh1*, and *Bdh2*. Empagliflozin also induced several oxidative transcripts, including *Acadl*, *Acadm*, and *Bdh1*, but did not suppress the lipogenic program. The concurrent induction of *Cyp4a10*, *Cyp4a14*, *Acot1*, and *Pdk4*, canonical PPARα targets that route fatty acids toward ω-oxidation and divert substrate flux away from glucose utilization, is consistent with the peroxisomal enrichment identified by gene set enrichment analysis and reinforces a coordinated program of hepatic metabolic remodeling with sotagliflozin. These transcriptional changes provide a hepatic correlate for the whole-body phenotype observed by indirect calorimetry, in which sotagliflozin-treated mice oxidized fat as their predominant substrate, and predict reduced hepatic *de novo* lipogenesis.

### Sotagliflozin enhances systemic β-oxidation and ketogenesis while suppressing hepatic *de novo* lipogenesis

Because circulating acylcarnitines reflect mitochondrial fatty acid oxidation efficiency, we profiled plasma acylcarnitines and free carnitine across all six groups. Vehicle-treated CKD mice accumulated long-chain species relative to sham controls, consistent with impaired β-oxidation in CKD (Figure 6A). Sotagliflozin reversed this signature, raising free carnitine and short- and medium-chain acylcarnitines while lowering long-chain species in both sham and CKD mice. Empagliflozin produced a directionally similar but weaker shift. Absolute plasma acylcarnitine concentrations are shown in supplemental table S3.

**Figure 6.**
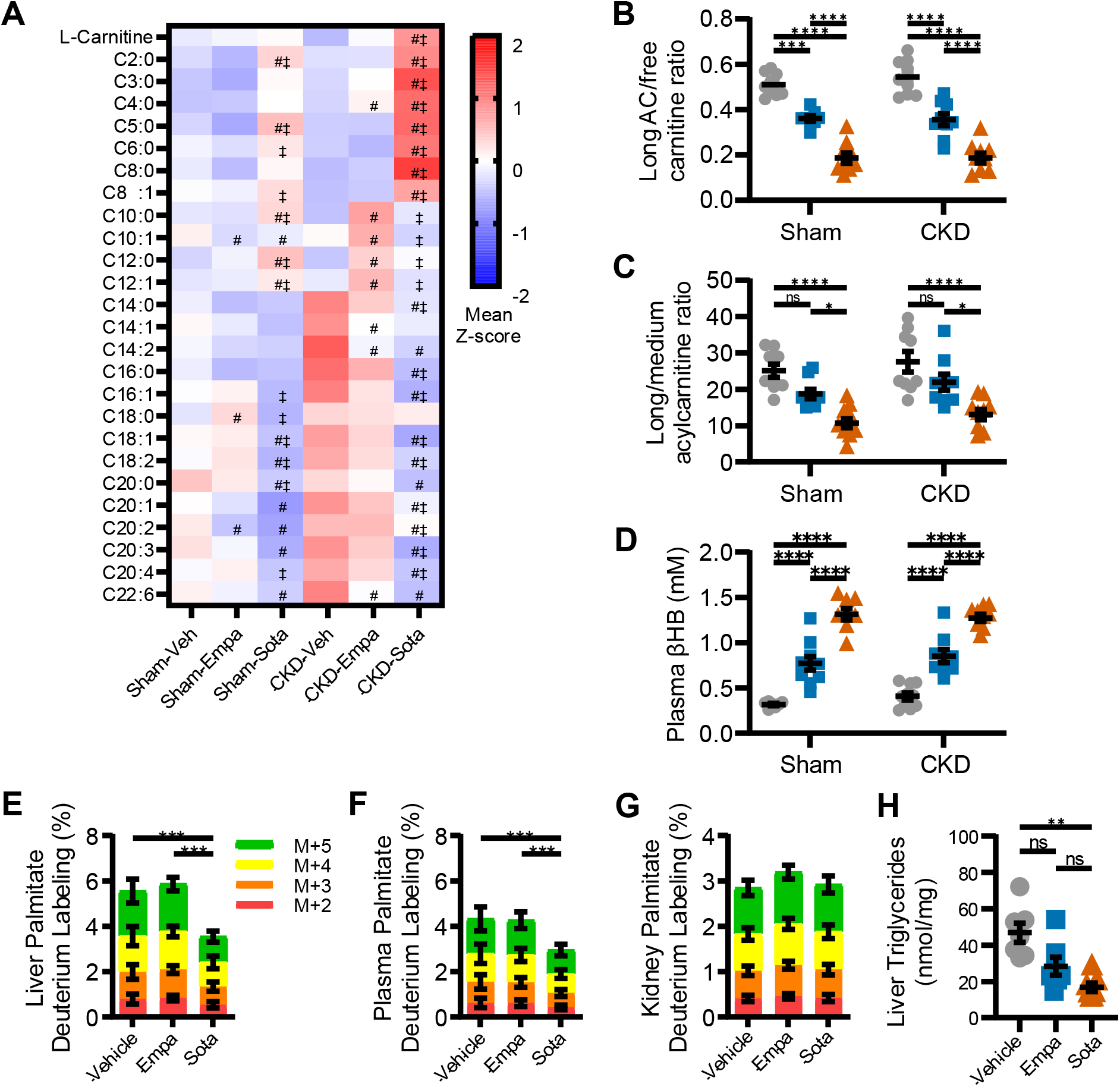
Sotagliflozin enhances systemic β-oxidation and ketogenesis while suppressing hepatic *de novo* lipogenesis. Plasma free carnitine and 25 AC species were quantified by targeted LC/MS in sham and CKD mice (n=9/group), and DNL was measured in vivo following ^2^H_2_O administration to isotopically label newly synthesized palmitate (n=9/group). (A) Mean standardized levels of plasma AC. Individual species were sum-normalized to the total AC pool within each sample and z-score standardized; cells display the group mean z-score. # indicates FDR *P*<0.05 versus vehicle; and ‡, FDR *P*<0.05 versus EMPA within the same surgical group.(B) Ratio of long-chain (C14-C22) AC to free carnitine absolute concentration. (C) Ratio of long-chain to medium-chain (C6-C12) AC absolute concentrations. (D) Plasma βHB. (E-G) Contributions of M+2, M+3, M+4, and M+5 isotopomers to total palmitate deuterium enrichment in liver (E), plasma (F), and kidney (G). Asterisks indicate significant differences in total percent enrichment between groups. (H) Hepatic triglyceride content normalized to tissue mass. Data are shown as mean±SEM. Comparisons in A through D were analyzed by two-way ANOVA with Tukey multiple comparisons test. Comparisons in E through H by one-way ANOVA. In B through H, ✱*P*<0.05, ✱✱*P*<0.01, ✱✱✱*P*<0.001, and ✱✱✱✱*P*<0.0001; ns, not significant. AC, acylcarnitine; βHB, β-hydroxybutyrate; CKD, chronic kidney disease; DNL, *de novo* lipogenesis; EMPA, empagliflozin; LC/MS, liquid chromatography-mass spectrometry; and SOTA, sotagliflozin.

Changes in acylcarnitine profile were also assessed by ratios. The ratio of long-chain to free carnitine, an index of carnitine sequestration by unmetabolized long-chain substrate, was reduced by both agents and reduced further by sotagliflozin in sham and CKD mice (Figure 6B). The ratio of long-chain to medium-chain acylcarnitines, which reports the completeness of chain shortening through β-oxidation, was significantly lowered only by sotagliflozin, with empagliflozin indistinguishable from vehicle (Figure 6C). Sustained β-oxidative flux also generates ketone bodies. Plasma β-hydroxybutyrate rose with both treatments and was highest in sotagliflozin-treated mice, regardless of surgical group (Figure 6D), consistent with the induction of *Hmgcs2*, *Bdh1*, and *Bdh2* observed in the liver (Figure 5G).

To determine whether increased fat oxidation was accompanied by reduced lipid synthesis, we measured *de novo* lipogenesis *in vivo* by deuterium labeling of newly synthesized palmitate. Sotagliflozin reduced palmitate deuterium enrichment in liver relative to both vehicle and empagliflozin (Figure 6E), and the same reduction was evident in plasma (Figure 6F), indicating lower hepatic output of newly synthesized fatty acid into the circulation. Kidney palmitate enrichment did not differ between groups (Figure 6G), localizing the lipogenic effect to the liver. Empagliflozin did not alter *de novo* lipogenesis in any compartment. This reduction in lipogenic flux was accompanied by lower hepatic lipid content: liver triglycerides were lower with sotagliflozin than with vehicle, whereas empagliflozin did not differ from vehicle (Figure 6H). Together, these data demonstrate that dual SGLT1/2 inhibition produces reciprocal changes in hepatic lipid handling, increasing β-oxidative flux and ketogenesis while simultaneously suppressing *de novo* lipogenesis, and that selective SGLT2 inhibition engages only the oxidative arm of this response.

## Discussion

Several studies have described mechanisms of cardiorenal protection by SGLT2 inhibitors that operate beyond glucosuria and glycemic control.^20–22^ Despite this progress, the cardiovascular benefit of the class remains curiously partitioned. Large outcome trials show consistent reductions in CKD progression, cardiovascular death, and heart failure hospitalization but little effect on myocardial infarction or stroke.^7,23^ Sotagliflozin, which inhibits both SGLT1 and SGLT2, is the only agent in the class to reduce atherothrombotic events in patients with CKD. However, SCORED enrolled only patients with diabetes and was terminated early for reasons unrelated to efficacy.^13,14^ No study has determined why dual inhibition produces this effect, or whether it extends to CKD without diabetes. Our study addressed this gap by examining whether additional SGLT1 blockade alters systemic substrate handling in a way that could account for atheroprotection.

To address these questions, we combined 5/6 nephrectomy with high-fat feeding in *Ldlr*^−/−^ mice to recapitulate features spanning the three domains of CKM syndrome in a single animal, including albuminuria, diastolic dysfunction, systemic inflammation, and insulin resistance. Under these conditions, sotagliflozin reduced atherosclerotic burden, adiposity, and systemic inflammation, whereas empagliflozin conferred only partial protection against atherosclerosis and no weight benefit. However, the two agents provided equivalent improvements in other cardiorenal metrics, including albuminuria and diastolic dysfunction, suggesting the divergence is metabolic rather than a difference in overall potency. While we were initially surprised that empagliflozin failed to alter body composition despite urinary caloric loss, this result aligns with prior work. Sweet Pee mice, which carry a loss-of-function mutation in *Slc5a2* encoding SGLT2, are glucosuric throughout life and are not protected against diet-induced obesity.^20,24^ Furthermore, in diet-induced obese mice, empagliflozin reduces body weight only with pair feeding, limiting compensatory hyperphagia.^25^ Loss of renal glucose reclamation therefore appears permissive rather than sufficient for cardiometabolic reprogramming.

We hypothesize that the cardiometabolic benefits of sotagliflozin stem from reduced and delayed prandial glucose availability. SGLT2 reclaims most filtered glucose in the proximal tubule, while SGLT1 absorbs dietary glucose across the intestinal brush border and recovers the residual filtered load in the S3 segment.^10^ Blocking SGLT2 alone diverts glucose already absorbed into the urine, leaving enteral absorption intact and allowing animals to compensate by increasing food intake, as shown in our empagliflozin-treated mice. Blocking both SGLT1 and SGLT2 simultaneously closes both entry and recovery pathways. Prandial glucose availability is therefore reduced and delayed regardless of how much the animal consumes, and a greater proportion of its energy demand must be met from lipid. Our calorimetry data support this, as only sotagliflozin sustained neutral energy balance and near-exclusive fat oxidation despite matched hyperphagia.

To determine whether this whole-body shift had a hepatic correlate, we profiled the liver transcriptome and measured lipogenic flux directly. SGLT2 inhibition is known to shift fuel selection toward lipid in humans and to engage hepatic oxidative programs in rodents^25,26^, and our gene set enrichment analysis reproduced this effect with empagliflozin. Additionally, our data reveal that dual SGLT1/2 inhibition significantly amplifies this phenotype while simultaneously suppressing hepatic *de novo* lipogenesis. This reciprocal shift is consistent with malonyl-CoA’s established role in regulating lipogenesis and β-oxidation via allosteric inhibition of carnitine palmitoyltransferase-1.^27,28^ That this occurred in mice consuming a high-fat diet and only under dual transporter blockade indicates that the operative signal is the absence of usable glucose due to SGLT1 inhibition rather than the abundance of dietary lipid.

Our data do not resolve how this metabolic reprogramming translates into reduced plaque, although several observations narrow the possibilities. SGLT1 is minimally expressed in vascular and immune cells, making direct action on the arterial wall unlikely. Furthermore, total plasma cholesterol was identical across treatment groups, arguing against a mechanism driven by total cholesterol exposure. The atheroprotection we observed is therefore likely to be remote and metabolic in origin. Sotagliflozin reduced hepatic *de novo* lipogenesis, which would be expected to lower VLDL-triglyceride secretion, consistent with the reduced plasma triglycerides and deuterium enrichment of plasma palmitate we observed. Triglyceride-rich lipoproteins and their remnants are causal drivers of atherosclerosis^29,30^, and sotagliflozin has been shown to lower postprandial triglycerides more than empagliflozin in patients with type 2 diabetes.^11^ Alternatively, the shift toward fatty acid oxidation and ketogenesis may directly alter plaque macrophage immunometabolism and phenotype. We have previously shown that CKD alters macrophage lipid metabolism in this model^15^ and that impaired β-oxidation accompanies the progression of CKM syndrome.^31,32^ Sotagliflozin reversed this signature and raised β-hydroxybutyrate, a ketone that suppresses NLRP3 inflammasome activation.^33^ The selective decrease in TNF-α with sotagliflozin, without changes in IL-6 or MCP-1, is consistent with a change in macrophage phenotype rather than broad suppression of inflammation. These mechanisms are not mutually exclusive, and both follow from the same underlying constraint on glucose availability.

Our findings also raise the possibility that dual SGLT1/2 inhibition is not merely an alternative within the SGLT2 inhibitor class but potentially a therapeutically superior approach for cardiometabolic protection. Several independent human observations support this rationale. Loss-of-function variants in *SLC5A1* are associated with a lower incidence of diabetes, heart failure, and cardiovascular death.^12^ In addition, across the SGLT2 inhibitor class, lower pharmacologic selectivity for SGLT2 over SGLT1 is associated with greater reductions in major adverse cardiovascular events.^34^ Empagliflozin and sotagliflozin lie at opposite ends of the SGLT2:SGLT1 selectivity spectrum, with ratios of 2500:1 and 20:1, respectively.^34^ Therefore, the presence of matched glucosuria results in significant differences in intestinal SGLT1 inhibition. In direct comparison with empagliflozin in patients with type 2 diabetes, sotagliflozin produced greater reductions in postprandial glucose and triglycerides and greater elevations in GLP-1.^11^ Our data suggest that the metabolic advantage of sotagliflozin arises specifically from constraining glucose availability at both absorptive and resorptive sites. If this applies to humans, patients with CKM syndrome and elevated atherothrombotic risk may benefit more from dual inhibition than from the SGLT2-selective agents that currently dominate prescribing.

The contribution of GLP-1 to the cardiometabolic phenotype observed with sotagliflozin is less clear. Intestinal SGLT1 blockade delivers unabsorbed glucose to L cells in the distal small intestine and stimulates endogenous GLP-1 release.^35^ While this increase in GLP-1 plausibly contributes to improved glucose tolerance, it is a weak candidate for atheroprotection, given that dipeptidyl peptidase-4 inhibitors raise endogenous GLP-1 to a comparable degree and are neutral for major adverse cardiovascular events.^36^ In contrast, the atherothrombotic benefit of GLP-1 receptor agonists is obtained at supraphysiologic GLP-1 receptor occupancy.^8^ A 3-fold rise in endogenous hormone is unlikely to reproduce what pharmacologic agonism achieves. Resolving this will require direct interrogation, either by GLP-1 receptor antagonism with exendin(9-39) or by intestine-specific deletion of *Slc5a1*. Whether sotagliflozin and a GLP-1 receptor agonist act additively is a separate question with immediate clinical relevance, particularly since combining a selective SGLT2 inhibitor with a GLP-1 receptor agonist confers no incremental cardiovascular benefit.^9^

There are limitations of our study. We used only male mice because cardiorenal protection by SGLT2 inhibitors is sex-independent in humans and female mice resist diet-induced atherosclerosis, but we have not excluded sex differences in intestinal SGLT1 expression. We matched doses by fractional excretion of glucose, the best available *in vivo* index of renal SGLT2 engagement. Since sotagliflozin also inhibits SGLT1 in the S3 segment, we cannot exclude a modest difference in SGLT2 blockade. However, we administered both agents at doses far exceeding clinical doses, producing maximal renal SGLT2 occupancy. Because sotagliflozin-treated mice weighed less than their comparators, we cannot fully separate weight-dependent and weight-independent contributions to atheroprotection. Treatment remained a significant determinant of plaque burden after ANCOVA adjustment for body mass (Figure 2I), but formal separation would require pair-feeding or weight-matched comparisons. Lastly, we calculated energy balance as the difference between ingested energy and measured expenditure and did not account for unabsorbed dietary carbohydrates. Because intestinal SGLT1 inhibition delivers glucose to the distal bowel, a portion of ingested energy may have been lost in stool or diverted to colonic fermentation into short-chain fatty acids. While this does not affect the measured increase in energy expenditure or the reduction in RER, unabsorbed dietary glucose could contribute to the negative energy balance and reduced adiposity observed with sotagliflozin. Future studies should determine whether the atheroprotection we observed persists with GLP-1 receptor blockade, whether it is reproduced by intestine-specific SGLT1 deletion, and whether sotagliflozin combined with a GLP-1 receptor agonist confers additive benefit.

In summary, sotagliflozin reciprocally reprograms hepatic lipid metabolism toward fatty acid oxidation and ketogenesis, and this shift is associated with greater atheroprotection than selective SGLT2 inhibition at equivalent glucosuria. These data support a model in which intestinal, not renal, glucose blockade drives cardiometabolic reprogramming in CKM syndrome.

## Data availability statement

RNA sequencing data are available in NCBI GEO (accession GSE345155). Analysis code and all other data supporting the findings of this study are available from the corresponding author upon reasonable request.

## Sources of Funding

This work was supported in part by U54DK137314, P30DK89503, and Juvenile Diabetes Research Foundation (JDRF) grant 5-COE-2019-861-S-B to SP. KS was supported by the KUHR training network U2C DK129445 and TL1DK136046 and by the AHA 26CDA1601970. VA Merit (I01CX002684-01), NIH R61/R33 (1R61HL177474-01), and the Mathers Foundation grant (AAL).

## Disclosures

The authors report no conflicts of interest to disclose.

## Supplemental Materials

Supplemental Methods

Figures S1-S2

Tables S1–S5

## Non-standard Abbreviations and Acronyms

2H2O: deuterated water
AC: acylcarnitine
ANCOVA: analysis of covariance
BUN: blood urea nitrogen
CKD: chronic kidney disease
CKM: cardiovascular-kidney-metabolic
CVD: cardiovascular disease
DNL: *de novo* lipogenesis
DPP-IV: dipeptidyl peptidase-4
E/A: ratio of early to late mitral inflow velocity
E/e′: ratio of early mitral inflow velocity to early diastolic mitral annular velocity
EMPA: empagliflozin
GLP-1: glucagon-like peptide-1
GSEA: gene set enrichment analysis
HFD: high-fat diet
IVRT: isovolumic relaxation time
LC/MS: liquid chromatography-mass spectrometry
Ldlr: low-density lipoprotein receptor
LV: left ventricular
MCP-1: monocyte chemoattractant protein-1
OGTT: oral glucose tolerance test
PCA: principal component analysis
PREVENT: Predicting Risk of cardiovascular disease EVENTs
Q-TOF: quadrupole time-of-flight
RER: respiratory exchange ratio
SGLT1: sodium-glucose cotransporter-1
SGLT2: sodium-glucose cotransporter-2
SOTA: sotagliflozin
TNF-α: tumor necrosis factor alpha
UACR: urine albumin-to-creatinine ratio
VCO2: carbon dioxide production
VO2: oxygen consumption
βHB: β-hydroxybutyrate

## Notes

### Competing Interest Statement

The authors have declared no competing interest.

