## Supplemental Material for "Sotagliflozin Reprograms Hepatic Lipid Metabolism and Attenuates Atherosclerosis Beyond Empagliflozin in CKM Mice"

### Supplemental Methods and Materials

#### Supplemental Methods

##### Animals

All animal experiments were performed in accordance with the approval of the University of Michigan Institutional Animal Care and Use Committee (IACUC) protocol. At 6 weeks of age, male *Ldlr*<sup>-/-</sup> mice (B6.129S7-Ldlr<sup>tm1Her/J</sup>, The Jackson Laboratory) were subjected to either sham operation or 5/6 nephrectomy to induce CKD, as previously described.<sup>1</sup> Briefly, the right kidney was removed in an initial procedure, followed by approximately two-thirds of the left kidney one week later. At 10 weeks of age, mice were transitioned to a high-fat diet (HFD; 20% protein, 21% fat with 0.21% cholesterol, and 50% carbohydrates by weight; D12079B, Research Diets Inc.) to induce hypercholesterolemia and block randomized to receive sotagliflozin (SOTA, 30mg/kg/day), empagliflozin (EMPA, 20mg/kg/day), or vehicle continuously via drinking water (n=20 per treatment arm within each surgical group, 2-3 mice/cage). SOTA and EMPA (Med Chem Express, Cat. #s HY-15516 and HY-15409) were dissolved in 100% ethanol with sonication, then diluted in water containing 0.25% Tween-80 to a final ethanol concentration of 1%. Vehicle-treated mice received drinking water containing 1% ethanol and 0.25% Tween-80. We selected final SOTA and EMPA concentrations in a preliminary dose-finding cohort, in which we calculated fractional glucose excretion from paired plasma and urine glucose and creatinine measurements after 7 days of treatment. Blood and urine glucose levels were measured using a handheld glucometer (AimStrip Plus, Germaine Laboratories Inc.) and an enzymatic assay (Abcam cat#ab65333), respectively. Plasma creatinine was measured by highly specific electrospray ionization and tandem LC-MS, as previously described.<sup>2</sup> The same measurement was repeated in the main cohort to confirm sustained pharmacodynamic activity and comparable renal SGLT2 inhibition between groups. All mice received HFD and drinking water ad libitum for 16 weeks before blood and tissue collection. Drinking water was refreshed every 48 hours, and body weight and food intake were measured weekly. Because studies in humans indicate that cardiorenal protection by SGLT2 inhibitors is independent of sex, and female mice are less susceptible to diet-induced atherosclerosis than males, only male mice were used for this study.<sup>3-5</sup>

##### **Oral glucose tolerance testing**

After 12 weeks of treatment, oral glucose tolerance testing (OGTT) was performed in sham and CKD mice (n = 5/group). Following a 6-hour fast, baseline blood glucose was measured by tail nick, and a 20% (w/v) dextrose solution was administered by oral gavage at 2 g/kg body weight (10 μL/g). Tail blood glucose was measured at 15, 30, 60, and 120 min post-gavage. In a separate subset of CKD mice (n = 5-6/group), plasma total GLP-1 levels were measured after OGTT by collecting blood in DPP-IV inhibitor (Millipore Sigma, Cat# DPDP4). Total GLP-1 levels were measured by ELISA (Millipore Sigma, Cat#EZGLP1T). Glycemic excursion and GLP-1 secretion were quantified as the total area under the curve.

##### **Metabolic Phenotyping**

At week 14, a subset of treated mice (n = 5 per group) in each surgical group underwent indirect calorimetry to measure energy expenditure. Mice were acclimated to single housing for 1 week before being transferred to Promethion cages (Comprehensive, High-Resolution Behavioral Analysis Systems, Sable Systems International, USA) at the University of Michigan Animal Phenotyping Core. Food and water consumption, x, y, and z beam breaks, VO<sub>2</sub>, and VCO<sub>2</sub> were measured at 20-minute intervals for 72 hours at 22°C. Animals were provided HFD and treatment water throughout the experiment. Respiratory exchange ratio (RER) was calculated as VCO<sub>2</sub>/VO<sub>2</sub>. Body composition was measured using an EchoMRITM-500 Body Composition Analyzer immediately before and after the 72 hr experiment, and average values were used for statistical analysis. Data were exported without reference to group identity, statistically compared by blinded core staff, and analyzed using ANCOVA with total body mass as the covariate in CalR (https://calrapp.org/). Data were then exported and visualized per International Indirect Calorimetry Consensus Committee (IICCC) guidelines.<sup>6</sup>

##### **Echocardiography measurements**

We performed transthoracic echocardiography after 15 weeks of treatment to assess changes in cardiac function. All echocardiographic measurements were recorded in anesthetized mice maintained with 1-2% isoflurane and 95% oxygen to maintain a heart rate of $450 \pm 50$ beats per minute. Briefly, transthoracic echocardiography was conducted using the Vevo F2 Imaging System (#53699-20, FUJIFILM Visualsonics, Inc., USA). The anterior chest hair was removed using Nair hair-removal cream before sedation. Prewarmed ultrasound gel was applied to the ventral thorax after the mice were immobilized on the stage. The parasternal short-axis view was used to obtain M-mode images for ejection fraction, fractional shortening, left ventricular mass, cardiac output, and other cardiac parameters. The apical four-chamber view was used for tissue Doppler and mitral valve pulse-wave Doppler measurements. A blinded analyzer measured all parameters at least 3 times; mean values were plotted for statistical analysis.

##### **Plasma Collection and Processing of the Heart and Aorta**

Mice were sedated with isoflurane, and blood was collected from the retro-orbital sinus into lithium heparin-coated microcapillary tubes. Mice were then humanely euthanized. An aliquot of blood was used to measure blood chemistries, electrolytes, hematocrit, hemoglobin, and BUN with an i-STAT-1 point-of-care analyzer (Abbott Laboratories). Additional plasma aliquots were flash-frozen in liquid nitrogen and stored at $-80^{\circ}\text{C}$ until use. The mouse was then perfused with sterile saline injected into the left ventricle. Hearts were separated from the aorta under a dissecting microscope and serially dehydrated in 15-30% sucrose before embedding in optimal cutting temperature compound. Aortas were extracted, fixed in 4% PFA for 24 hours, then transferred to sterile saline for storage and subsequent processing.

##### **Plasma analyte measurements**

Plasma creatinine levels were measured by highly specific electrospray ionization and tandem LC/MS, as previously described.<sup>2</sup> Plasma levels of TNF-α, IL-10, IL-6, MCP-1, insulin, and Cystatin C were measured by a custom ELISA using a MESO QuickPlex SQ 120MM analyzer (UPLEX, Meso Scale Discovery) per manufacturer's instructions. Plasma total cholesterol and triglycerides were quantified by colorimetric assays (Fujifilm Wako) per manufacturer's instructions.

##### ***En face* atherosclerosis quantification**

*En face* atherosclerosis was performed and quantified, as described previously.<sup>7</sup> Briefly, aortas were removed from storage in sterile saline, cleaned free of adventitia, and stained with Oil Red O to visualize neutral lipids. The aorta was then cut open along the outer curvature down to the subclavian artery, and the inner curvature was then cut open from the ascending aorta down to the iliac bifurcation. The aorta was pinned flat with dissection pins onto a tray fitted with Sudan-black-stained paraffin. Images were taken on a Zeiss Stemi 2000-C dissecting microscope with a 48MP camera. Atherosclerotic lesions were quantified using ImageJ Fiji software and presented as percent lesion area. All quantification was verified by 2 blinded observers.

##### **Aortic sinus atherosclerosis quantification**

Atherosclerotic lesions in the aortic sinus were sectioned, stained, and analyzed as described previously.<sup>7</sup> Briefly, hearts were thawed from -80°C storage in optimal cutting temperature and cut parallel to the tricuspid valve approximately 3 mm from the base of the ventricles. The aortic sinus was placed perpendicular to the bottom of a tissue mold and embedded in optimal-cutting-temperature compound. At the appearance of the tricuspid valve leaflets, the aortic sinus was serially sectioned in 8-μm increments over 8 microscope slides with 3 sections per slide. Sections were then stained with H&E, mounted, and lesion area quantified using ImageJ Fiji software. Total lesion area was summed across serial slides, and quantifications were verified by 2 blinded observers.

##### ***In vivo* assessment of *de novo* lipogenesis**

To assess relative differences in *de novo* lipogenesis between SOTA and EMPA-treated mice, we administered deuterated water ($^2\text{H}_2\text{O}$) to isotopically label newly synthesized palmitic acid, the primary product of fatty acid synthase, with $^2\text{H}$. Mice were given intraperitoneal boluses of $^2\text{H}_2\text{O}$ (20 μl/gm body weight) 24 and 72 h prior to tissue collection. Lipids were extracted from plasma, kidney, and liver samples using a modified Bligh and Dyer method with water:methanol:dichloromethane (2:2:2). The organic layer containing lipids was collected, hydrolyzed to liberate palmitate, and analyzed by LC/MS. *De novo* lipogenesis was measured as a function of percentage $^2\text{H}$ enrichment of palmitate, as previously described, using an Agilent 6546 Q-TOF mass spectrometer, coupled to an Agilent 1290 LC.<sup>1</sup> The acquired raw data were processed using Mass Hunter Profinder (Agilent Profinder V10). The percent labeling of palmitate was quantified by measuring the peak area of each mass shift relative to unlabeled palmitate. Total palmitate enrichment in $^2\text{H}$ was determined from mass isotopomers (M+2, M+3, M+4, and M+5). Each sample was measured in triplicate, and the average was used as the sample value.

##### **Plasma acylcarnitine profiling**

Acylcarnitines (AC) play a crucial role in transporting long-chain fatty acids into the mitochondria, and circulating levels reflect the mitochondrial AC pool and the efficiency of β-

oxidation. Enhanced mitochondrial function induced by SGLT2 inhibitors has been associated with increased plasma levels of short- (C2-C5) and medium-chain (C6-C12) acylcarnitines without changes in long-chain (C14-C22) acylcarnitines, supporting improved fatty acid flux through β-oxidation.<sup>8</sup> We quantified plasma acylcarnitines (n=25) and free carnitine in sham- and CKD-treated mice by LC/MS using an Agilent 6490 Triple Quadrupole tandem mass spectrometer with a targeted method described previously.<sup>9</sup> Internal standards are listed in (supplemental Table S1). Raw mass spectrometry signals for each AC were normalized to the corresponding authenticated internal standard for absolute quantification. Within each sample, individual ACs were sum-normalized as a fraction of total AC, capturing relative class-level abundance and avoiding misleading interpretations that arise when comparing absolute concentrations between samples with differing total AC pools. Sum-normalized values were z-score standardized to allow side-by-side comparison in heatmaps, since absolute AC concentrations span $\mu\text{mol/L}$ to $\text{nmol/L}$. For each standardized acylcarnitine species, group differences were assessed by two-way ANOVA with Tukey's post hoc test within each surgical group, and the resulting P values for the sotagliflozin-versus-vehicle and sotagliflozin-versus-empagliflozin contrasts were corrected for the 26-analyte panel using the Benjamini-Hochberg procedure ($\text{FDR} < 0.05$). For quality assurance and quality control, we ran a reference pool at the beginning and after every 10 samples to assess reproducibility and drift in mass spectrometry performance over time. Coefficient of variation for each acylcarnitine analyte are shown in supplemental Table S2.

##### **Liver triglyceride quantification**

Total liver triglyceride content was quantified in flash-frozen liver samples by colorimetric assay (Abcam, Cat# ab65336) per the manufacturer's instructions at 1:20 dilution and normalized to tissue mass.

#### **RNA sequencing**

Total RNA was extracted from liver tissue of CKD mice (n=6/group) using PureLink RNA Mini Kit (Thermo Fisher), following the manufacturer's instructions. Plasmidsaurus performed RNA library preparation with 3' end tagging, incorporating unique molecular identifiers (UMIs) for PCR and optical duplicate removal. After quality control, sequencing was performed on an Illumina NovaSeq X Plus, producing 10 million single-end 90 bp reads per sample. Reads were aligned to the *Mus musculus* GRCm39 reference genome with STAR v2.8.1a, deduplicated with `umi_tools` v1.1.6, and quantified to gene-level counts with `featureCounts` (Rsubread v2.12.3). Genes with $\geq 10$ counts in $\geq 3$ samples and annotated as protein-coding or lncRNA were retained. Differential expression was assessed with DESeq2 using median-of-ratios normalization and a negative binomial model with treatment group as the sole factor; pairwise Wald contrasts were computed for sotagliflozin vs. vehicle, empagliflozin vs. vehicle, and sotagliflozin vs. empagliflozin, with log2 fold changes shrunk using the `ashr` estimator and significance defined as Benjamini–Hochberg FDR $< 0.05$ and $|\log_2 \text{fold change}| \geq 1$. Variance-stabilizing transformed counts were used for principal component analysis and for per-gene Z-scored heatmaps. Gene set enrichment analysis was performed with `fgsea` on genes ranked by the Wald statistic against the MSigDB mouse Hallmark collections (n=50 sets), yielding a normalized enrichment score (NES) and Benjamini-Hochberg-adjusted P value for each gene set. Over-representation analysis of differentially expressed gene lists was performed separately using the hypergeometric test, yielding an adjusted P value for each gene set.

#### **Statistics**

Data are presented as mean±SEM. Normality was assessed by the Shapiro-Wilk. Variables that deviated from normality were log-transformed before analysis or compared by nonparametric tests, as indicated in the corresponding figure legend. Measurements obtained in both surgical groups were analyzed by two-way ANOVA with surgical group (sham or CKD) and treatment (vehicle, EMPA, or SOTA) as fixed factors, followed by Tukey's multiple comparisons test. Measurements obtained only in CKD animals were analyzed by one-way ANOVA with treatment group as a fixed factor. Animals that died before the study ended were excluded from terminal endpoints. Serial measurements, including body weight, blood glucose, and total GLP-1 during OGTT, were analyzed by two-way repeated-measures ANOVA. Glycemic and incretin excursions were additionally summarized as total AUC by the trapezoidal method and compared as above. A two-sided $P < 0.05$ was considered statistically significant. Analyses were performed using GraphPad Prism version 11 (GraphPad Software) and R version 4.6.1 (R Foundation for Statistical Computing).

**Figure S1: Serum Chemistries in Sham- and CKD-treated Mice**

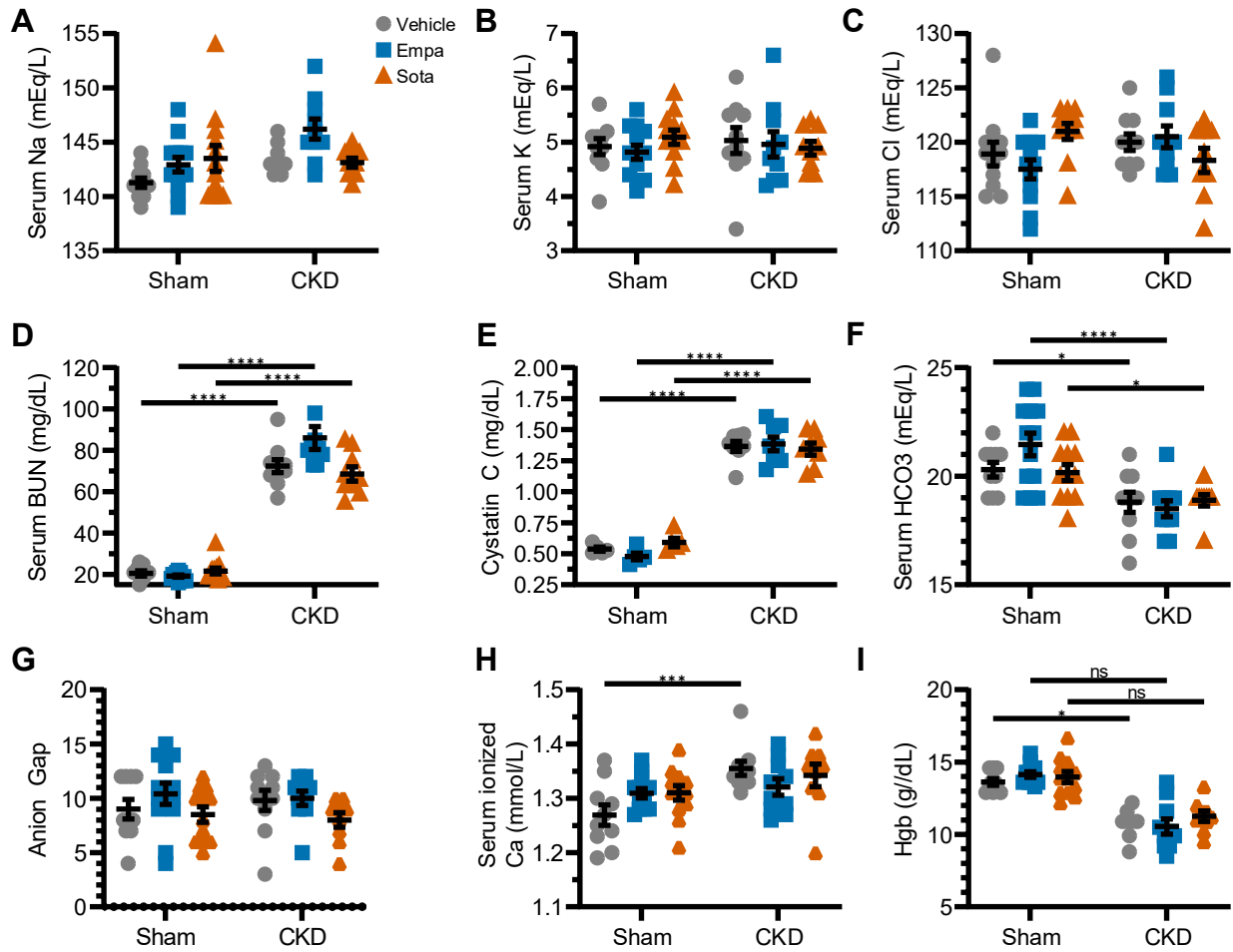

**Figure S1. Serum chemistries confirm chronic kidney disease without treatment-related electrolyte or acid-base disturbance.** Blood was collected at the time of tissue harvest after 16 weeks of treatment. Electrolytes, BUN, bicarbonate, and hemoglobin were measured on an i-STAT-1 point-of-care analyzer and plasma cystatin C by multiplex immunoassay ($n=10-12$/group). (A) Serum sodium. (B) Serum potassium. (C) Serum chloride. (D) BUN. (E) Plasma cystatin C. (F) Serum bicarbonate. (G) Anion gap. (H) Serum ionized calcium. (I) Hemoglobin. Data are shown as mean ± SEM. Comparisons were analyzed by two-way ANOVA with Tukey multiple comparisons test. \*$P < 0.05$, \*\*$P < 0.01$, \*\*\*$P < 0.001$, and \*\*\*\*$P < 0.0001$; ns, not significant. BUN, blood urea nitrogen; CKD, chronic kidney disease; EMPA, empagliflozin; Hgb, hemoglobin; and SOTA, sotagliflozin.

**Figure S2: Indirect Calorimetry Results for Sham-treated Mice**

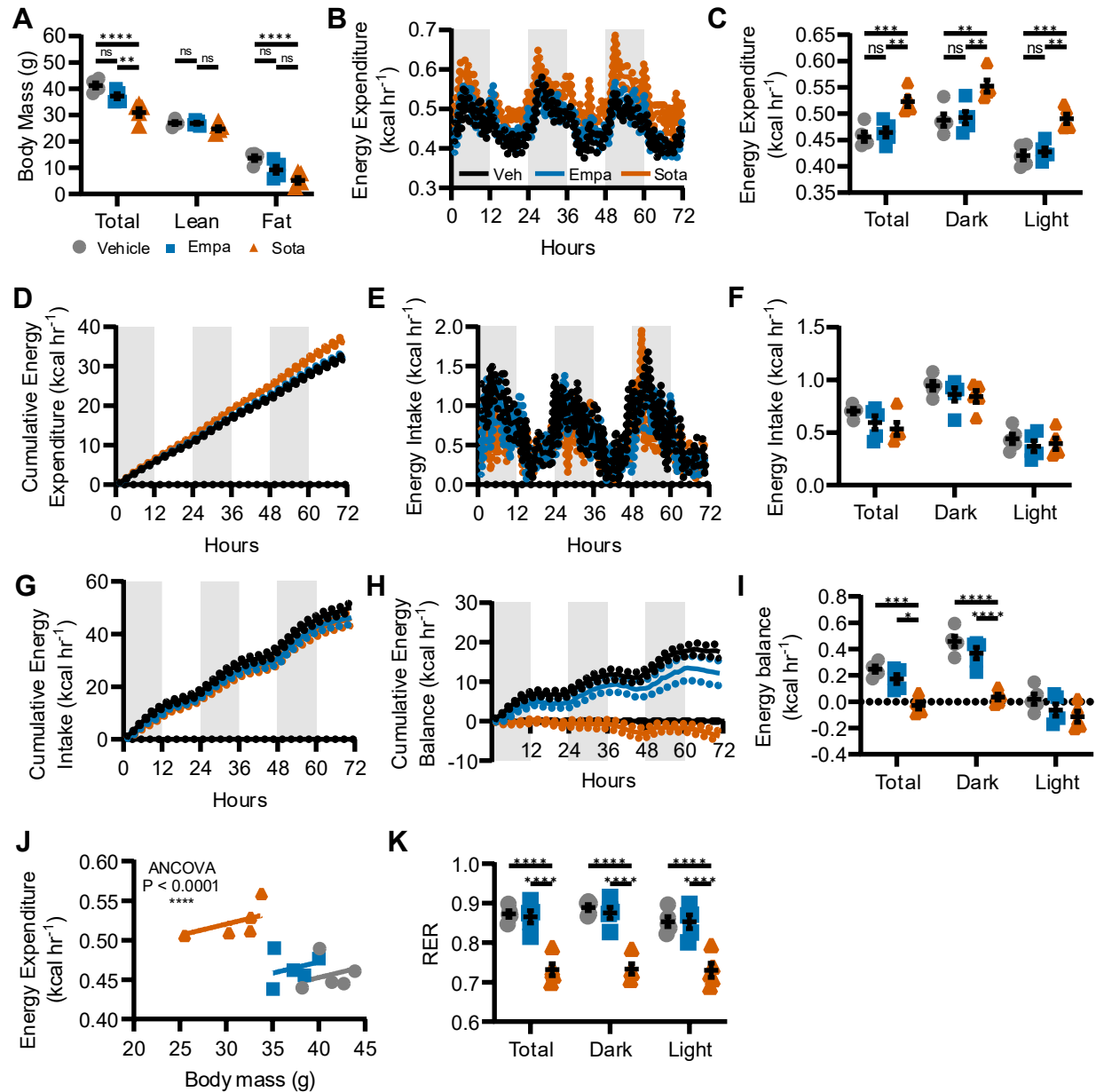

**Figure S2. Sotagliflozin increases energy expenditure and shifts substrate utilization toward fatty acid oxidation in sham mice.** Sham mice underwent indirect calorimetry for 72 hours after 14 weeks of treatment (n=5/group), with body composition measured by EchoMRI immediately before and after the recording. (A) Total, lean, and fat mass. (B-C) Energy expenditure over the 72-hour recording (B) and averaged across the total recording and by dark and light phase (C). (D) Cumulative energy expenditure. (E-F) Energy intake over the recording period (E) and averaged across the total recording and by phase (F). (G) Cumulative energy intake. (H-I) Cumulative energy balance (H) and energy balance averaged across the total recording and by phase (I). (J) ANCOVA of energy expenditure against total body mass. (K) RER, calculated as VCO<sub>2</sub>/VO<sub>2</sub>. Shaded regions in B, D, E, G, and H denote the dark phase; solid lines

indicate group means and dotted lines indicate SEM. Data are shown as mean±SEM. Energy expenditure was analyzed by ANCOVA against total body mass in CalR and is presented per IICCC guidelines; all other comparisons were analyzed by one-way ANOVA with Tukey multiple comparisons test. \*$P<0.05$, \*\*$P<0.01$, \*\*\*$P<0.001$, and \*\*\*\*$P<0.0001$; ns, not significant. EMPA, empagliflozin; RER, respiratory exchange ratio; SOTA, sotagliflozin.

**Table S1:** List of authenticated internal standards for acylcarnitine measurement

| Component (Isotopic Label, Enrichment) | Purity | Gravimetric Target Concentration (uM) | Concentration by Gravimetry (uM) | Analyzed Concentration (uM) |
| --- | --- | --- | --- | --- |
| L-Carnitine (N-trimethyl-D<sub>9</sub>, 98%) | 99.7% | 152.0 | 157.5 | 152.1 ± 13.3 |
| L-Acetylcarnitine (N-methyl-D<sub>3</sub>, 98%) | 99.5% | 38.0 | 38.1 | 37.4 ± 2.7 |
| L-Propionylcarnitine (N-N-methyl-D<sub>3</sub>, 98%) | 98.1% | 7.6 | 7.6 | 7.8 ± 0.6 |
| L-Butyrylcarnitine (N-methyl-D<sub>3</sub>, 98%) | 98.7% | 7.6 | 7.6 | 7.4 ± 0.2 |
| L-Isovalerylcarnitine (N-trimethyl-D<sub>9</sub>, 98%) | >98% | 7.6 | 7.6 | 7.5 ± 0.2 |
| L-Octanoylcarnitine (N-methyl-D<sub>3</sub>, 98%) | 99.5% | 7.6 | 7.6 | 7.5 ± 0.2 |
| L-Myristoylcarnitine (N-trimethyl-D<sub>9</sub>, 98%) | >98% | 7.6 | 7.6 | 7.2 ± 0.3 |
| L-Palmitoylcarnitine (N-methyl-D<sub>3</sub>, 98%) | >98% | 15.2 | 15.2 | 14.9 ± 0.5 |

**Table S2:** Measured carnitines and acylcarnitines and their coefficient of variation

| Class | Sub-class | Compound by carbon number and number of double bonds in acyl chain | Coefficient of variation (%) |
| --- | --- | --- | --- |
| Carnitines | Carnitine | L-Carnitine | 11.79 |
|  | Acylcarnitines (AC) | C(2:0) | 1.41 |
|  |  | C(3:0) | 0.69 |
|  |  | C(4:0) | 5.68 |
|  |  | C(5:0) | 7.23 |
|  |  | C(6:0) | 9.99 |
|  |  | C(8:1) | 3.10 |
|  |  | C(8:0) | 2.82 |
|  |  | C(10:1) | 6.68 |
|  |  | C(10:0) | 2.91 |
|  |  | C(12:1) | 5.82 |
|  |  | C(12:0) | 7.29 |
|  |  | C(14:1) | 40.58 |
|  |  | C(14:0) | 5.60 |
|  |  | C(14:2) | 55.73 |
|  |  | C(16:1) | 22.76 |
|  |  | C(16:0) | 7.73 |
|  |  | C(18:2) | 26.46 |
|  |  | C(18:1) | 4.24 |
|  |  | C(18:0) | 3.93 |
|  |  | C(20:4) | 35.69 |
|  |  | C(20:3) | 14.65 |
|  |  | C(20:2) | 20.64 |
|  |  | C(20:1) | 36.11 |
|  |  | C(20:0) | 14.10 |
|  |  | C(22:6) | 60.76 |

**Table S3:** Absolute acylcarnitine concentrations by CKD status and treatment

| Analyte (μM) | Sham Mean (95% CI) |  |  | CKD Mean (95% CI) |  |  |
| --- | --- | --- | --- | --- | --- | --- |
|  | Vehicle | Empa | Sota | Vehicle | Empa | Sota |
| Carnitine | 620.2<br>(368.1-872.3) | 662.7<br>(529.9-795.5) | 690.9<br>(522.1-859.7) | 515.8<br>(357.6-673.9) | 672.2<br>(507.32-837.0) | 936.3<br>(567.6-1305.1) |
| AC2:0 | 889.0<br>(740.5-1037.4) | 858.5<br>(680.9-1036.1) | 1128.9<br>(890.6-1367.1) | 928.7<br>(756.5-1100.8) | 934.1<br>(708.2-1160.0) | 1325.7<br>(964.4-1687.0) |
| AC3:0 | 18.89<br>(14.68-23.09) | 16.44<br>(13.78-19.09) | 23.33<br>(15.70-30.96) | 19.83<br>(16.11-23.56) | 22.74<br>(18.25-27.23) | 35.98<br>(27.55-44.41) |
| AC4:0 | 12.14<br>(9.88-14.39) | 12.34<br>(8.59-16.08) | 16.16<br>(8.20-24.12) | 13.22<br>(9.26-17.18) | 17.03<br>(11.23-22.82) | 26.12<br>(19.73-32.50) |
| AC5:0 | 1.73<br>(1.35-2.12) | 1.466<br>(1.08-1.85) | 2.54<br>(1.90-3.18) | 1.68<br>(1.36-2.01) | 1.67<br>(0.80-2.54) | 3.18<br>(2.58-3.80) |
| AC6:0 | 0.650<br>(0.402-0.898) | 0.510<br>(0.119-0.901) | 0.768<br>(0.496-1.040) | 0.468<br>(0.373-0.564) | 0.383<br>(0.185-0.580) | 1.211<br>(0.594-1.829) |
| AC8:0 | 0.361<br>(0.232-0.490) | 0.282<br>(0.193-0.372) | 0.403<br>(0.297-0.509) | 0.279<br>(0.238-0.320) | 0.308<br>(0.191-0.425) | 0.705<br>(0.529-0.881) |
| AC8:1 | 0.013<br>(0.008-0.019) | 0.013<br>(0.008-0.017) | 0.015<br>(0.012-0.018) | 0.010<br>(0.005-0.015) | 0.011<br>(0.008-0.014) | 0.018<br>(0.013-0.023) |
| AC10:0 | 0.480<br>(0.271-0.690) | 0.524<br>(0.345-0.702) | 0.671<br>(0.444-0.899) | 0.416<br>(0.305-0.528) | 0.806<br>(0.420-1.191) | 0.506<br>(0.349-0.663) |
| AC10:1 | 0.967<br>(0.375-2.309) | 0.402<br>(0.227-0.578) | 0.588<br>(0.205-0.971) | 0.796<br>(0.388-1.203) | 1.468<br>(0.055-2.991) | 0.369<br>(0.186-0.551) |
| AC12:0 | 3.436<br>(2.228-4.644) | 4.758<br>(3.115-6.402) | 6.550<br>(2.436-10.663) | 3.419<br>(2.675-4.164) | 6.106<br>(3.671-8.540) | 4.750<br>(3.692-5.808) |
| AC12:1 | 1.429<br>(0.984-1.874) | 1.727<br>(1.225-2.230) | 1.970<br>(1.017-2.924) | 1.593<br>(1.082-2.103) | 2.376<br>(0.961-3.790) | 1.521<br>(1.209-1.834) |
| AC14:0 | 10.61<br>(6.73-14.50) | 7.66<br>(5.15-10.18) | 8.48<br>(6.23-10.73) | 15.69<br>(10.91-20.48) | 12.77<br>(7.73-17.82) | 9.59<br>(7.93-11.24) |
| AC14:1 | 4.66<br>(3.42-5.901) | 3.84<br>(2.41-5.26) | 3.48<br>(2.57-4.38) | 5.84<br>(4.40-7.28) | 4.32<br>(3.01-5.63) | 4.07<br>(2.91-5.23) |
| AC14:2 | 0.181<br>(0.136-0.226) | 0.148<br>(0.107-0.189) | 0.149<br>(0.110-0.189) | 0.284<br>(0.198-0.371) | 0.172<br>(0.134-0.210) | 0.150<br>(0.097-0.204) |
| AC16:0 | 42.22<br>(28.52-55.91) | 31.90<br>(25.78-37.94) | 34.40<br>(28.27-40.53) | 62.00<br>(46.66-77.32) | 53.03<br>(41.59-64.46) | 33.70<br>(27.21-40.18) |
| AC16:1 | 17.97<br>(12.14-23.81) | 16.87<br>(6.98-26.75) | 12.49<br>(9.95-15.03) | 25.60<br>(18.77-32.44) | 19.06<br>(15.63-22.48) | 11.85<br>(9.20-14.50) |
| AC18:0 | 20.84<br>(15.28-26.40) | 24.82<br>(3.89-45.76) | 14.92<br>(10.87-18.96) | 26.87<br>(19.33-34.42) | 25.85<br>(21.31-30.39) | 24.65<br>(17.39-31.92) |
| AC18:1 | 32.05<br>(22.94-41.17) | 30.42<br>(13.61-47.23) | 23.47<br>(17.18-29.76) | 40.33<br>(30.44-50.21) | 34.43<br>(28.14-40.72) | 20.53<br>(16.06-24.99) |
| AC18:2 | 3.197<br>(2.338-4.057) | 3.234<br>(1.060-5.409) | 2.190<br>(1.570-2.810) | 4.153<br>(3.022-5.283) | 3.536<br>(2.822-4.250) | 2.548<br>(1.950-3.146) |
| AC20:0 | 1.839<br>(0.450-3.227) | 1.231<br>(0.023-2.439) | 0.602<br>(0.254-0.950) | 1.511<br>(0.246-2.776) | 1.188<br>(0.622-1.754) | 0.489<br>(0.135-0.842) |
| AC20:1 | 2.917<br>(2.434-3.400) | 2.273<br>(1.156-3.389) | 1.787<br>(1.420-2.153) | 3.706<br>(2.697-4.715) | 3.250<br>(2.541-3.958) | 2.575<br>(1.704-3.447) |
| AC20:2 | 0.227<br>(0.104-0.349) | 0.147<br>(0.098-0.196) | 0.128<br>(0.094-0.162) | 0.250<br>(0.178-0.323) | 0.251<br>(0.211-0.291) | 0.201<br>(0.147-0.254) |
| AC20:3 | 0.046<br>(0.029-0.064) | 0.034<br>(0.014-0.054) | 0.024<br>(0.016-0.031) | 0.057<br>(0.040-0.074) | 0.047<br>(0.031-0.063) | 0.024<br>(0.016-0.032) |
| AC20:4 | 0.812<br>(0.553-1.072) | 0.758<br>(0.224-1.292) | 0.603<br>(0.411-0.795) | 1.040<br>(0.756-1.324) | 0.895<br>(0.738-1.052) | 0.599<br>(0.495-0.704) |
| AC22:6 | 0.408<br>(0.303-0.513) | 0.324<br>(0.198-0.449) | 0.292<br>(0.210-0.375) | 0.546<br>(0.347-0.744) | 0.361<br>(0.241-0.481) | 0.271<br>(0.144-0.398) |

**Table S4:** Differentially expressed genes in sotagliflozin-treated mice compared to vehicle and empagliflozin

| Gene Symbol | Gene Name | Sotagliflozin vs Vehicle |  | Sotagliflozin vs Empagliflozin |  |
| --- | --- | --- | --- | --- | --- |
|  |  | Log2 FC | Adjusted P | Log2 FC | Adjusted P |
| *Hsd17b6* | hydroxysteroid (17-beta) dehydrogenase 6 | 3.36 | 4.31E-48 | 1.02 | 4.14E-05 |
| *Tff3* | trefoil factor 3, intestinal | -5.37 | 2.59E-22 | -2.55 | 4.61E-05 |
| *Fasn* | fatty acid synthase | -3.4 | 8.45E-17 | -2.1 | 8.50E-08 |
| *Mmp12* | matrix metalloproteinase 12 | 4.44 | 1.26E-15 | 2.42 | 1.65E-05 |
| *Gpnmb* | glycoprotein (transmembrane) nmb | 4.98 | 1.19E-14 | 2.83 | 1.65E-05 |
| *Bdh1* | 3-hydroxybutyrate dehydrogenase 1 | 1.66 | 2.78E-07 | 2.06 | 2.60E-04 |
| *Ccl4* | C-C motif chemokine ligand 4 | 2.29 | 3.19E-07 | 1.47 | 1.10E-03 |
| *Col1a1* | collagen, type I, alpha 1 | 1.32 | 3.13E-06 | 1.3 | 5.36E-05 |
| *Sqle* | squalene epoxidase | -2.84 | 3.87E-06 | -2.31 | 3.59E-04 |
| *Acadl* | acyl-Coenzyme A dehydrogenase long chain | 1.15 | 8.17E-06 | 2.28 | 4.61E-04 |
| *Trem2* | triggering receptor expressed on myeloid cells 2 | 1.44 | 1.19E-05 | 1.52 | 6.88E-05 |
| *Acadm* | acyl-Coenzyme A dehydrogenase medium chain | 1.26 | 1.49E-05 | 1.5 | 1.48E-03 |
| *Atf3* | activating transcription factor 3 | 1.65 | 1.81E-05 | 1.39 | 9.35E-04 |
| *Srebf1* | sterol regulatory element binding transcription factor 1 | -2.18 | 1.47E-04 | -2.74 | 9.58E-03 |
| *Acacb* | acetyl-CoA carboxylase beta | -1.52 | 2.52E-03 | -1.1 | 1.43E-05 |

Genes differentially expressed in sotagliflozin-treated mice versus both vehicle and empagliflozin (n = 6 per group). Significance was defined as Benjamini-Hochberg adjusted $P < 0.05$ and $|\log_2 \text{fold change}| \geq 1$ in each contrast, tested independently. A positive $\log_2$ fold change indicates higher expression with sotagliflozin. FC indicates fold change.

**Table S5:** Number of animals analyzed for each experiment and assay

| Completed study | Sham Veh | Sham Empa | Sham Sota | CKD Veh | CKD Empa | CKD Sota |
| --- | --- | --- | --- | --- | --- | --- |
| Body wt | 12 | 13 | 13 | 20 | 14 | 16 |
| OGTT | 5 | 5 | 5 | 5 | 5 | 6 |
| GLP-1 | — | — | — | 5 | 5 | 6 |
| Calor. | 5 | 5 | 5 | 6 | 5 | 5 |
| Echo | 12 | 12 | 13 | 12 | 10 | 10 |
| En face | 12 | 13 | 13 | 14 | 14 | 14 |
| Sinus | 12 | 13 | 13 | 14 | 14 | 14 |
| Chem. | 11 | 13 | 13 | 11 | 14 | 14 |
| Cytokines | 8 | 8 | 8 | 8 | 8 | 8 |
| AC panel | 9 | 9 | 9 | 9 | 9 | 9 |
| <sup>2</sup>H flux | 5 | 5 | 5 | — | — | — |
| Liver TG | — | — | — | 10 | 10 | 10 |
| RNA-seq | — | — | — | 6 | 6 | 6 |
